# Few-shot learning of predictive features with dendrites and behavioural timescale synaptic plasticity in the hippocampus

**DOI:** 10.64898/2026.05.08.723802

**Authors:** Colleen J. Gillon, Claudia Clopath

## Abstract

When an animal enters a new environment, neurons in the hippocampus begin to map out the space. They become selectively responsive to features like the animal’s location, the position of rewards, and the presence of stimuli relevant to navigating or performing a specific task. With experience, hippocampal neurons also develop behaviour-related biases. It is thus believed that the hippocampus encodes multi-sensory, behaviourally-relevant cognitive maps of environments that are critical to navigation, learning and guiding behaviour. The predictive learning hypothesis proposes that these complex maps emerge because a core goal of the brain is to learn to predict the features of its environment. In sensory cortex, predictive learning provides a compelling explanation of anticipatory and error-like sensory responses. Pyramidal neurons receive top-down and bottom-up inputs to their proximal basal and distal apical dendrites, respectively. These complementary inputs streams are thought to enable them to act as comparison units, signaling discrepancies between predictive and sensory inputs. In the hippocampus, however, the potential link between pyramidal neurons and predictive learning is still underexplored. Here, we investigate the possibility that pyramidal neurons perform a similar comparator function in the hippocampus. In our model, two-compartment pyramidal neurons receive sensory information about salient features of the environment at their distal apical dendrites which is compared to tuned spatial inputs received more proximally to their cell body. We demonstrate how predictive learning implemented in this circuit using behavioural timescale synaptic plasticity and distal apical inhibition can explain a variety of spatial and behaviourally-relevant features encoded in the hippocampus. We also lay out key predictions for validating our model experimentally. As such, our work helps bridge an important gap in the literature on predictive learning in the hippocampus and set the stage for more robust experimental validation of this prominent hypothesis.

## 1 Introduction

As we interact with the world, we are flooded with streams of sensory information. Our brains must parse this information quickly and effectively to productively guide our behaviour. A leading theory in computational neuroscience of how the brain does this is the predictive learning hypothesis. According to this hypothesis, instead of interpreting stimuli from scratch at each moment, the brain learns an internal model of the world. This model allows the brain not only to anticipate expected or familiar stimuli in the environment, but also to rapidly recognise and respond to unexpected or novel information. Under this hypothesis, when unpredicted stimuli are encountered, the brain incorporates this novel information, updating its internal model of the world to better predict future sensory inputs [Srinivasan et al., 1982; Rao & Ballard, 1999; Huang & Rao, 2011; Bastos et al., 2012; Keller & Mrsic-Flogel, 2018; Millidge et al., 2022; Aizenbud et al., 2026]. Research on sensory processing in the brain provides compelling support for the predictive learning hypothesis. In humans and animals, unexpected stimuli elicit strong cortical responses that cannot be explained by their sensory properties alone [Näätänen et al., 1978; Garrido et al., 2009; Wacongne et al., 2011]. Within individual brain regions, and particularly in sensory areas, pyramidal neurons have been shown, through experience, to develop not only error responses [Musall et al., 2017; Jordan & Keller, 2020; Audette et al., 2022; Gillon et al., 2024], but also predictive responses, firing in anticipation of their preferred stimulus in a familiar context [Fiser et al., 2016; Leinweber et al., 2017; Garrett et al., 2020; Audette et al., 2022; Zhou & Schneider, 2024]. Together, this evidence supports the idea that a core goal of the brain is to learn to predict its environment.

Although relatively simple and immediate sensory predictions may be computed locally in sensory brain areas, predictions reflecting more abstract and behaviourally-determined relationships between stimuli and involving longer time dependencies likely require the input of higher-order brain areas [Keller & Mrsic-Flogel, 2018]. During spatial navigation, for example, effectively predicting upcoming sensory inputs requires an understanding of an environment’s structure. The hippocampus, which plays a critical role in memory formation and learning the structure of new environments, has thus emerged as a strong candidate for more abstract, behaviourally-dependent predictive learning [Kumaran & Maguire, 2006; Exton-McGuinness et al., 2015; Fiser et al., 2016; Doron et al., 2020; Sinclair et al., 2021; George et al., 2023a; Aquino et al., 2024; Spens & Burgess, 2024]. It is well established that when an animal enters a new environment, hippocampal neurons begin to encode features of the environment. Most notable are place cells, often studied in areas CA3 and CA1, which respond selectively to specific locations in an environment [O’Keefe & Dostrovsky, 1971; O’Keefe & Conway, 1978]. Hippocampal neurons, particularly in area CA1, have also been shown to respond selectively to task-relevant features like reward-related features [Hollup et al., 2001; Lee et al., 2012; Gauthier & Tank, 2018; Ormond & O’Keefe, 2022; Terada et al., 2022; Yun et al., 2023; Qian et al., 2025; Sosa et al., 2025; Bausch et al., 2026; Yaghoubi et al., 2026], cues and other sensory features [Alexander et al., 2020; Keinath et al., 2020; Moore et al., 2021a; Radvansky et al., 2021; Sheridan et al., 2024], temporal features [MacDonald et al., 2011], and conjunctions of these features [Komorowski et al., 2009; Alexander et al., 2020; Xiao et al., 2020; Omer et al., 2022]. The hippocampus is thought to encode these features to form abstract cognitive maps that enable an animal to effectively navigate and interact with both new and familiar environments [O’Keefe & Dostrovsky, 1971; O’Keefe & Conway, 1978; Morris et al., 1982; Breese et al., 1989; Muller & Kubie, 1989; Mehta et al., 1997; McNaughton et al., 2006; Smith & Mizumori, 2006; Eichenbaum & Cohen, 2014; Robinson et al., 2020; Kim et al., 2026].

Computational neuroscientists have postulated that the hippocampus’ role in extracting and encoding task-relevant features emerges from its role in predictive learning [Levy, 1989; Hasselmo & Schnell, 1994; Lynch & Granger, 1992; Blum & Abbott, 1996; Kumaran & Maguire, 2007; Lisman & Redish, 2009; Stachenfeld et al., 2017; Santos-Pata et al., 2021; Chen et al., 2024; Levenstein et al., 2024; Bennett et al., 2025; Liu et al., 2025]. However, how feature learning through predictive learning might be implemented in the hippocampus at the circuitry level has not been well explored computationally. In sensory processing research, numerous computational studies have demonstrated how pyramidal neurons, the principal neurons in the cortex, are uniquely suited to supporting predictive learning [Urbanczik & Senn, 2014; Guerguiev et al., 2017; Sacramento et al., 2018; Hertäg & Sprekeler, 2020; Payeur et al., 2021; Mikulasch et al., 2023], as they receive top-down and bottom-up inputs at their proximal basal and distal apical dendrites, respectively [Marques et al., 2018] and integrate these nonlinearly via large events like dendritic spikes and plateau potentials [Larkum et al., 2001; Hay et al., 2016]. Notably, closely analogous properties are observed in hippocampal circuitry. Pyramidal neurons in areas CA1 and CA3 receive indirect inputs from entorhinal cortex layer 2 (EC2) via the perforant pathway and direct inputs from layer 3 (EC3) via the temporoammonic pathway. The indirect inputs, transmitted via the dentate gyrus and area CA3, target dendrites more proximal to the cell body and convey spatial information derived, at least in part, from the activity of grid cells and other spatially-modulated neurons in entorhinal cortex [Klausberger & Somogyi, 2008; Moser et al., 2008; Barry & Burgess, 2014; Lee et al., 2020]. The direct inputs from EC3 target the distal apical dendrites of pyramidal neurons, and have been proposed to transmit target-like information about salient features of the environment [Vago & Kesner, 2008; Deshmukh & Knierim, 2011; Grienberger & Magee, 2022; Bowler & Losonczy, 2023; Issa et al., 2024; Dorian et al., 2026]. As in sensory cortex, coincident input to both compartments can generate strong depolarisation events, including plateau potentials which are sustained periods of high neural activity lasting hundreds of milliseconds. Remarkably, plateau potentials have been shown to trigger behavioural timescale synaptic plasticity (BTSP), a type of plasticity that operates over multiple seconds and induces large, one-shot synaptic changes sufficient for a robust new place field to emerge in a single behavioural trial [Bittner et al., 2015, 2017; Milstein et al., 2021; O’Hare et al., 2022; Gonzalez et al., 2024; O’Hare et al., 2025; Madar et al., 2025b; Dorian et al., 2026]. The asymmetry of the BTSP kernel in area CA1, which lends a slight anticipatory bias, and its broad timecourse point to a potential role in learning predictive environmental features [Cho & McClelland, 2025; Li et al., 2024]. In this paper, we demonstrate how few-shot predictive learning of the behaviourally-relevant features of a new environment can emerge directly from these key properties of the hippocampal circuit. To show this, we build a rate-based neural circuit centered on two-compartment CA1-like pyramidal neurons. Each pyramidal neuron receives spatial inputs at a proximal compartment and inputs signaling salient features of the environment at a distal compartment. As a simulated agent navigates a new environment, coordinated activity across both compartments drives plateau potentials when salient features are encountered. As demonstrated in the experimental literature, these plateau potentials can act as discrete events triggering BTSP-like plasticity at the radial oblique dendrite inputs, modeled here as the pyramidal neuron’s proximal compartment. Through BTSP-triggering plateau potentials, the CA1-like pyramidal neurons in our model learn to anticipate the salient features of the environment the simulated agent is navigating. Critically, each pyramidal neuron also inhibits its own distal compartment via an oriens lacunosum moleculare (OLM)-like distal apical dendrite targeting inhibitory interneuron [Geiller et al., 2020; Milstein et al., 2021; Topolnik & Tamboli, 2022; Campbell et al., 2025; Kipper et al., 2025; Udakis et al., 2025; Vaasjo et al., 2025]. This self-inhibition ensures each neuron can, through learning, reduce the frequency and intensity of plateau potentials, enabling the overall circuit to stabilise as increasingly predictive features are learned. We demonstrate, in a linear track simulation, that our model allows CA1-like pyramidal neurons to form a stable field around a salient feature, like a landmark object, and recapitulates known place field formation properties, like the linear relationship between running velocity and place field width [Bittner et al., 2017; Milstein et al., 2021; O’Hare et al., 2022; Rolotti et al., 2022; Gonzalez et al., 2024; O’Hare et al., 2025]. We then show that this same circuit, with only slightly adjusted parameters, also forms stable place fields around salient features during navigation of a much more complex two-dimensional environment. Importantly, through BTSP, these place fields are also stably updated when the structure of the environment changes through the addition of teleportation ports. By bringing together key insights from hippocampal, sensory and predictive learning research, our model provides a biologically-grounded proof of concept for how few-shot learning of predictive features may be implemented in the hippocampal circuitry. Furthermore, our simulations lay out testable predictions for future research, and our codebase provides a reusable framework for expanding the proposed circuit and generating new predictions.

## 2 Results

### 2.1 Hippocampal circuit model for few-shot place field formation through BTSP

To investigate the potential role of the hippocampal circuitry in predictive learning, we first aimed to build a circuit model in which a pyramidal neuron reliably formed a stable place field after a single internally triggered BTSP-like learning event. Using `RatInABox` [George et al., 2024], we designed a rate-based neural circuit comprising three neuronal areas: (1) a place cell area, broadly representing CA3, (2) an object cell area, broadly representing EC3, and (3) a pyramidal neuron area, broadly representing CA1 (see Fig. 1A). Activity was modeled for neurons in all three areas while a simulated agent traveled at around 0.25 m/s (±0.05 m/s) along a spatially continuous linear track, encountering a landmark object at the three-fifth mark (see Fig. 1B).

**Figure 1:**
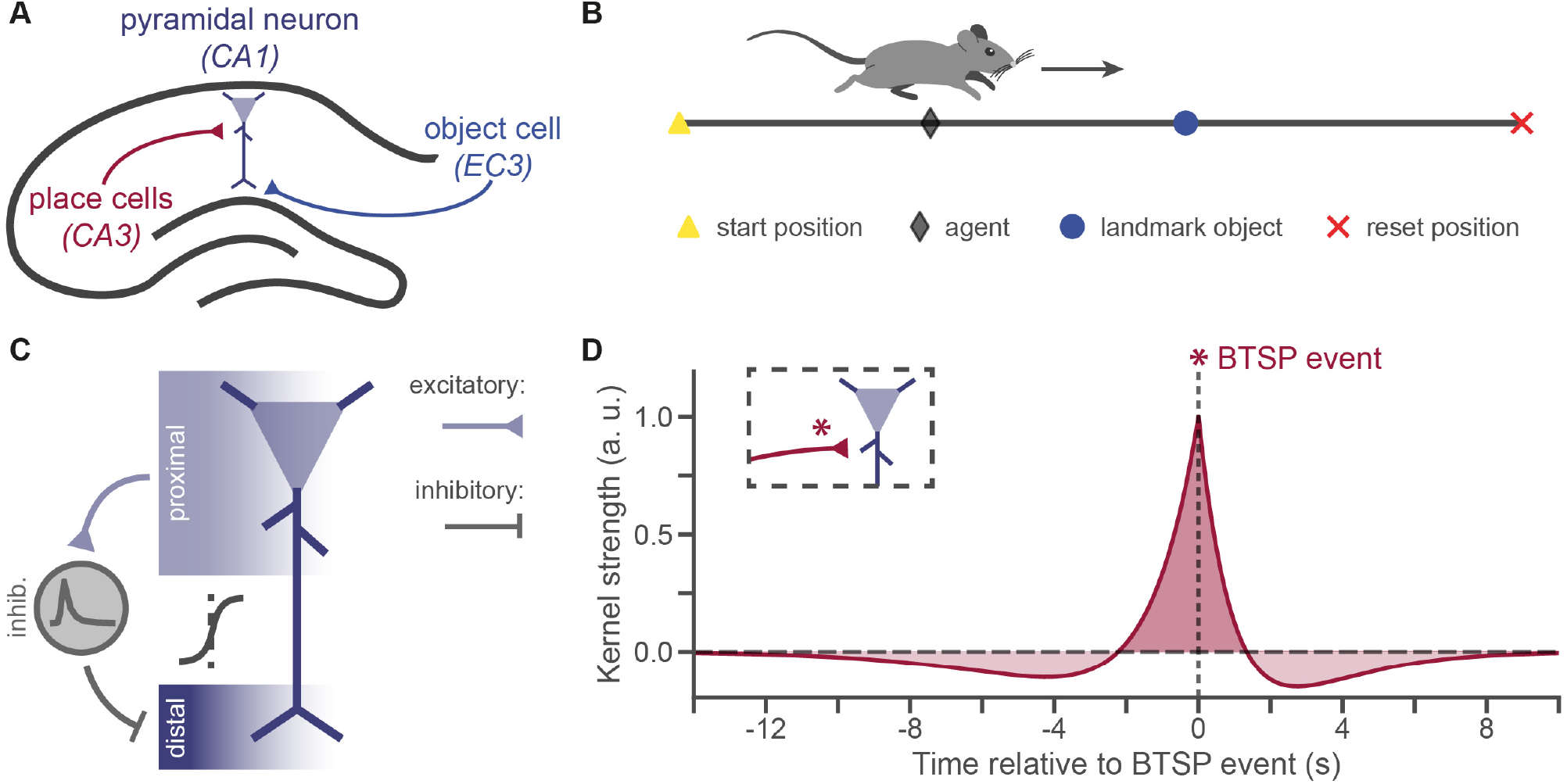
Circuit model of place field formation through BTSP in the hippocampus. **A**. Schematic of the hippocampal circuit. Place cells (CA3, red) send excitatory projections to the proximal compartment of a pyramidal neuron (CA1, purple), while an object cell (EC3, blue) sends excitatory projections to the pyramidal neuron’s distal compartment. **B**. Schematic of the linear track. For each run of the track (6 m), the agent (black diamond) begins at the start position (yellow triangle) and runs toward the reset position (red x), encountering a landmark object (blue circle) 3/5’s of the way down the track. The agent is shown here halfway to the landmark object, with a not-to-scale cartoon running mouse above it. An arrow shows its direction of travel. **C**. Schematic of the pyramidal neuron circuit (CA1). Each pyramidal neuron comprises two separate compartments: a proximal compartment (light purple, top) and a distal compartment (dark purple, bottom). The compartments are non-linearly connected, with each receiving a sigmoid-filtered input from the other. An inhibitory neuron paired to the pyramidal neuron receives excitatory inputs from the proximal compartment, which also serves as the output of the neuron. The interneuron filters this input, and in turn inhibits the pyramidal neuron’s distal compartment. **D**. Weight-update kernel used to induce BTSP-like plasticity at place cell synapses (CA3) targeting the pyramidal neuron (CA1). Broadly, weights for inputs activated within a few seconds (approx. -2 to 1.7 sec) of a BTSP-triggering plateau potential (BTSP event) are potentiated, while weights for inputs activated further away in time are depotentiated. Kernel strength is plotted in arbitrary units (a. u.) with the peak normalised to 1.0.

We modeled all neurons as single compartment neurons, except for pyramidal neurons which were two-compartment neurons (see Fig. 1C). In each simulation, the number of pyramidal neurons was matched to the number of landmark objects. Thus, in the linear track simulations, only one pyramidal neuron was modeled. The pyramidal neuron’s distal compartment represented the distal apical dendritic tuft, while its proximal compartment represented the cell body and the radial oblique dendrites. The basal dendrites were omitted for simplicity, as they show less spatial tuning diversity and do not appear to undergo BTSP [O’Hare et al., 2022; Gonzalez et al., 2024; Jain et al., 2024]. To drive strong activity in the distal compartment when the agent was near or at the landmark object, we provided narrow Gaussian input from an object cell, with a high, fixed weight. This connection roughly corresponds to the temporoammonic pathway from EC3 to CA1 in the hippocampus. In contrast, we targeted the proximal compartment with uniform and initially weak inputs from broader Gaussian place cells tiling the linear track. These inputs roughly correspond to the Schaffer collaterals projecting from CA3 to CA1 (see Fig. 1A). To enable plateau potential dynamics to emerge in the pyramidal neuron, we also simulated an NMDA receptor-mediated current in each compartment [Takahashi & Magee, 2009; Grienberger et al., 2014; Bittner et al., 2017].

All learning occurred in the weights of the place cell neurons targeting the proximal compartment of the pyramidal neuron. One-shot plasticity was triggered discretely each time a BTSP-triggering plateau potential, defined here as 100 ms or more of sustained high neural activity, occurred in the pyramidal neuron’s proximal compartment. Specifically, place cell weights onto the proximal compartment of the pyramidal neuron were updated following a BTSP-like learning rule [Milstein et al., 2021] (see Fig. 1D), with divisive weight normalisation applied only if the neuron’s total weights passed a specified threshold. Given the extended timecourse of the full BTSP kernel (see Fig. 1D), weight updates were always applied 20 simulated seconds after they had been triggered. This was the minimum time required to ensure that the full update could be computed and applied in a one-shot manner across all synapses involve. Remarkably, a very similar synaptic update delay has been observed experimentally, with the cellular changes underlying synaptic plasticity taking at least 20 seconds to take effect following a BTSP event [Jain et al., 2024]. During this short plasticity expression delay period, we temporarily blocked additional BTSP events. Finally, we allowed the pyramidal neuron to inhibit its own distal compartment via a paired interneuron receiving filtered inputs from the proximal compartment (see Fig. 1C). Through this delayed self-inhibition, the pyramidal neuron could block BTSP-triggering plateau potentials from continuing to occur after learning, and thus prevent runaway BTSP-driven weight updates. This critically enabled it to learn self-stabilising predictive fields.

### 2.2 Stable place fields form through one-shot learning on a linear track

Our first aim, as mentioned above, was to design a model that replicates the one or few-shot formation of place fields through BTSP that has been observed experimentally. This was achieved in our model by enabling a pyramidal neuron to develop a place field around a target object in the environment through BTSP, and then to stabilise the field by anticipating and inhibiting subsequent inputs to its distal compartment. We tested our model by simulating four full traversals of the linear track described above (see Fig. 2A). To simplify visual interpretation of the results, at the end of each traversal, the agent was held at the start point for 20 sec, allowing any plasticity resulting from a plateau potential that occurred during the previous track traversal to be applied before the next traversal began. As intended, during each track run, the object cell was strongly activated near its target, the landmark object, whereas place cell activity tiled the linear track uniformly (see Fig. 2A). BTSP-triggering plateau potentials consistently occurred on the first traversal, at the location of the object, and a place field was then formed through BTSP (see Fig. 2B). On subsequent traversals, the pyramidal neuron then expressed a place field around the object. The place field’s shape reflected our asymmetrical BTSP kernel (see Fig. 1D), and featured an initial spike resulting from the simulated NMDA receptor-mediated currents [Antic et al., 2010]. Importantly, its broad timecourse enabled the distal dendritic inhibition to ramp up sufficiently on each subsequent traversal such that input from the object cell to the distal compartment could no longer induce BTSP-triggering plateau potentials (see Fig. 2A, C). In other words, using a BTSP-like internally triggered plasticity rule, the pyramidal neuron in our circuit was able to stably form a place field around a target object in a one-shot manner.

**Figure 2:**
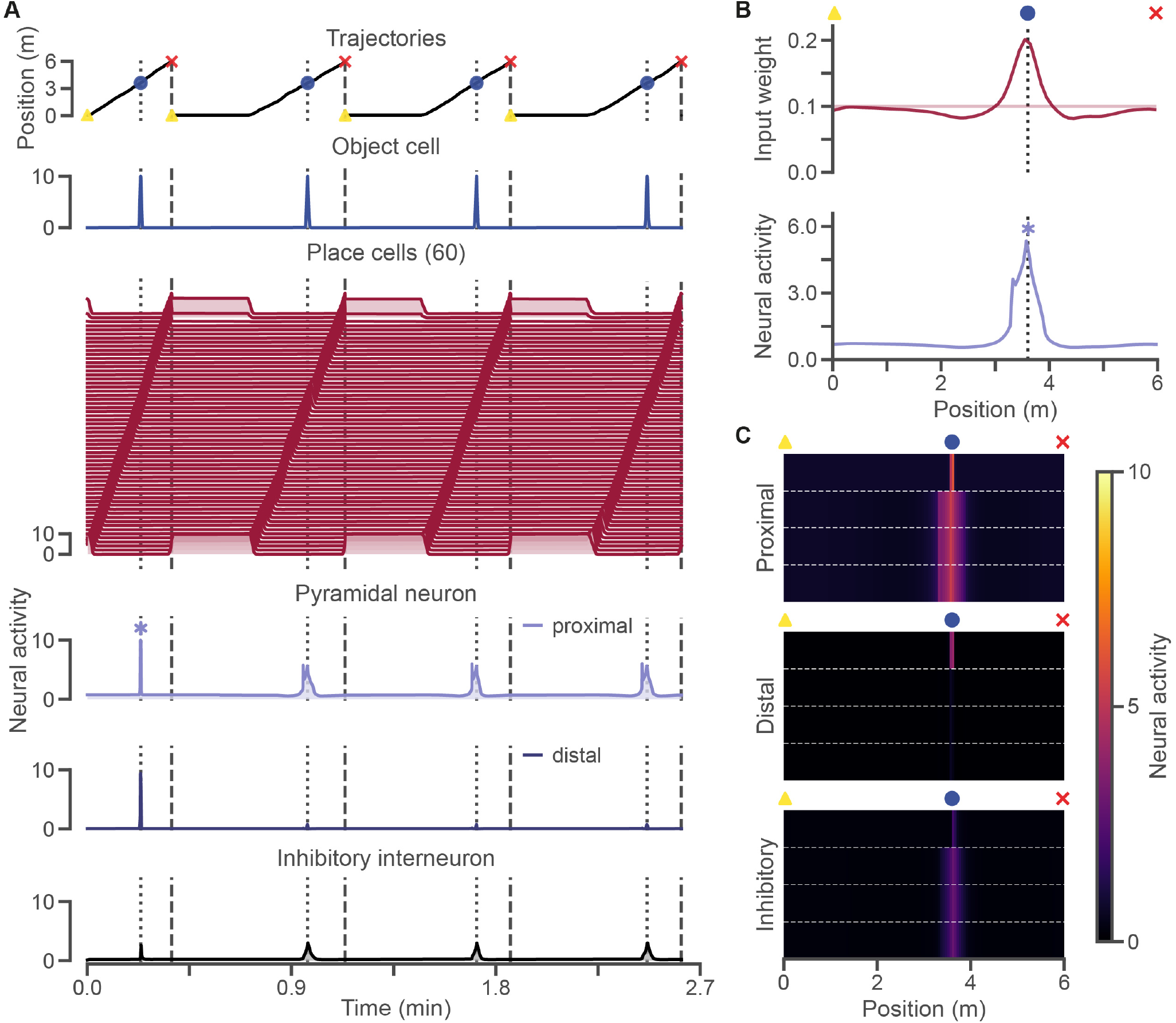
Place fields formed through BTSP on a linear track are self-stabilising. **A**. Agent position (*top*) across four track traversals with icons showing the start (yellow triangle), landmark object (blue circle) and reset positions (red x). Neural activity (*bottom*) across track traversals for the object cell (blue), all 60 place cells (red), and the three components of the pyramidal neuron circuit: the proximal compartment (light purple, *top*), the distal compartment (dark purple, *middle*) and the inhibitory interneuron (black, *bottom*). The x-axis shows time in simulated minutes. A BTSP-triggering plateau potential occurred during the first traversal only (light purple asterisk). Following this, the proximal compartment of the pyramidal neuron developed a place field around the object. This field was inherited and filtered by the inhibitory interneuron, suppressing distal activity around the object. Neural activity is expressed in arbitrary units, and ranges from 0 to 10 for each neuron or compartment. **B**. Following the BTSP event, place cell weights (*top*) formed an asymmetrical place field around the object, and the proximal compartment of the pyramidal neuron (*bottom*) expressed a place field around the object. The light purple asterisk marks the location where the BTSP-triggering plateau potential occurred. **C**. Spatially binned neural activity (120 bins) from **A** (*bottom*) in each of the three components of the pyramidal neuron circuit: the proximal compartment (*top*), the distal compartment (*middle*) and the inhibitory interneuron (*bottom*). Activity for the four track traversals is plotted from top to bottom, and separated by white, horizontal dotted lines.

The parameters of the model were specifically adjusted to enable the stable one-shot formation of a place field exemplified in Fig. 2. However, for each parameter and parameter combination tested, the network’s behaviour generally changed smoothly from being unable to undergo BTSP (i.e., no BTSP-triggering plateau potentials), to consistently stabilising after a single BTSP update (i.e., experiencing only one BTSP-triggering plateau potential), to being unable to stabilise even after several BTSP updates (i.e., experiencing numerous BTSP-triggering plateau potentials) (see Fig. S1 and S2). For the final model, presented here, parameter values lying in the middle of the range of values yielding the target behaviour (a single BTSP event) were retained (see Tables 1 and 2). Thus, the ability of the model to enable stable one-shot place field formation is not unique to the specific set of parameter values used, but instead shared across a range of parameter values.

Lastly, it should be noted that, in this simulation, the hippocampal circuit’s ability to completely abolish BTSP-triggering plateau potentials in the proximal compartment after initial learning is only possible because the environment and simulation conditions are highly stable. As demonstrated in later simulations, changes to the environment and variability in agent behaviour can lead to additional BTSP-triggering plateau potentials and new learning. Experimentally, it has been shown that plateau potentials, including BTSP-triggering ones, continue to occur in hippocampal neurons after initial learning [Milstein et al., 2021; O’Hare et al., 2025]. We propose that this is due to two main factors: on-going incremental plasticity mechanisms not modeled here, and real world experimental conditions which are much more complex and changeable, and thus not perfectly predictable. The aim of our model, however, is not to fully explain neural activity in the hippocampus, but rather to demonstrate how our self-stabilising learning rule can theoretically enable the hippocampus to learn a perfectly predictive map of the environment and abolish BTSP-triggering plateau potentials.

### 2.3 Place field width correlates with running velocity

Having shown that our circuit could form a stable place field on a linear track through a one-shot BTSP learning event, we next asked whether the resulting place fields recapitulated known properties of *in vivo* place fields formed through BTSP. Specifically, we asked whether place field width was linearly correlated to the running velocity of the agent during the BTSP event, as has been observed experimentally [Bittner et al., 2017; Milstein et al., 2021; O’Hare et al., 2022; Rolotti et al., 2022; Gonzalez et al., 2024; O’Hare et al., 2025]. We ran simulations in which the agent ran at different velocities during the BTSP induction stage, ranging from 0.05 to 0.40 m/s. Once a stable place field was formed, evaluation traversals were simulated at a highly variable speed (0.25 ± 0.25 m/s) to obtain reliable and comparable estimates of the resulting place fields. We estimated place fields for each running velocity by averaging neural activity in the proximal compartment across the evaluation traversals and computed the width as full-width at half maximum after smoothing.

In most cases, the place field formed during the induction stage was robust enough to block additional BTSP-triggering plateau potentials in the evaluation phase. However, in a few low speed cases (3/29), the initial place field was too narrow to provide sufficient distal inhibition at the higher evaluation phase speeds, and an additional BTSP event or two occurred. In these cases, weight normalisation was recruited, preventing runaway weight updates. Place fields were computed only from traversals that occurred after the final weight update was applied. Given that our BTSP kernel is applied in a time-dependent manner, we expected that place field width would closely reflect the agent’s speed as this directly determined how much of the track the agent traversed in the seconds before and after reaching the landmark object. As predicted, our model was able to recapitulate the relationship observed experimentally, namely that place field width increased linearly with running velocity (see Fig. 3). In the context of predictive learning, this suggests that the parameters of the hippocampus’ first encounter of a feature in an environment provides a strong inductive bias, with the circuit assuming that the feature will generally be encountered under similar circumstances, e.g., at a similar speed [Pecirno & Keinath, 2025]. As observed above, this inductive bias appeared to be generally adequate, as the initial place fields formed in our simulations tended to generalise well to later encounters of the landmark object at different speeds during the evaluation phase. The exception to this was the very narrow place fields formed at the lowest running velocities. These did not generalise well to higher speeds, with additional plateau potentials and plasticity being triggered during the evaluation phase. Thus, BTSP learning allowed the circuit not only to form an initial hypothesis about how the landmark object would typically be encountered, but also to then correct it if it proved inadequate.

**Figure 3:**
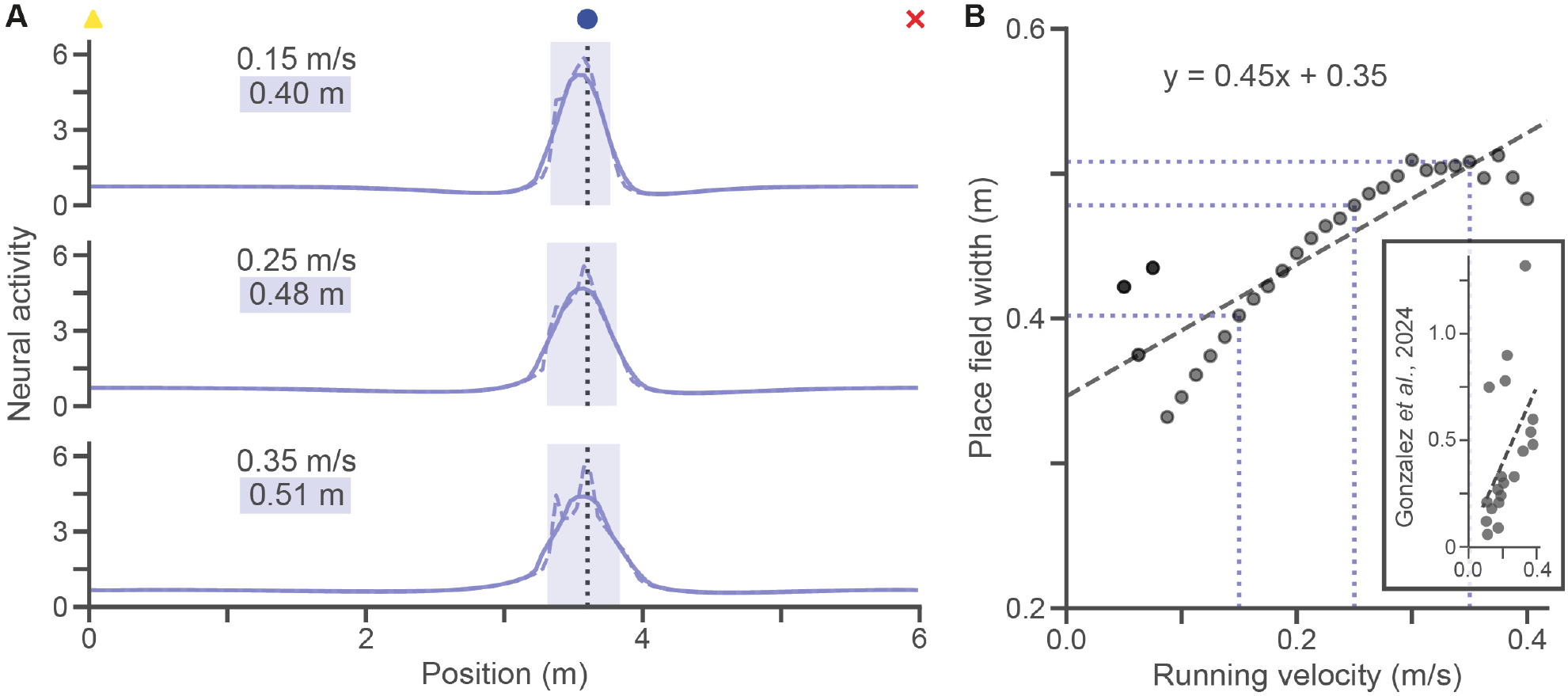
Place field width is linearly correlated to the agent’s running velocity when BTSP was triggered. **A**. Examples of place fields formed at different running velocities: 0.15 m/s (*top*), 0.25 m/s (*middle*), and 0.35 m/s (*bottom*). Icons at the top show the start (yellow triangle), landmark object (blue circle), and reset (red x) positions. The vertical dashed line also shows the object location. Place fields are computed by averaging neural activity across several track traversals following the last BTSP update. Dotted lines show averaged neural activity, while full lines show the smoothed neural activity used to robustly compute place field widths: 0.40 m (*top*), 0.48 m (*middle*), and 0.51 m (*bottom*). **B**. Relationship between running velocity (m/s) and place field width (m). Place field width tends to increase linearly with the agent’s running velocity during the BTSP event. Data points for which a second (1/29) or third (2/29) BTSP event occurred are marked by increasingly darker dots. The dashed line shows a linear regression (*y* = 0.45*x* + 0.35). Examples from **A** are marked with light purple dotted lines. See Fig. S3 for the same results expressed in terms of input weights. The inset reproduces Gonzalez et al. [2024]’s experimental results showing a linear relationship between running velocity and place field width.

It should also be noted that in our simulations at higher speeds (*>* 0.35 m/s), place field widths were consistently lower than predicted based on running velocity. Since the width of the place cell weights continued to increase linearly at these speeds (see Fig. S3), this effect is likely due to nonlinear dynamics specific to our pyramidal neuron model. At high speeds, BTSP learning in our model produces weaker weights, reducing the ability of pyramidal neurons to enter a high neural activity regime. This, in turn, disproportionately narrows the place fields computed for these higher speeds. Accordingly, at the lower speeds, when additional BTSP events occurred, this greatly increased place field width without much effect on place cell weight widths (see Fig. 3 and S3). In real brains, this effect may be compensated for by on-going incremental plasticity or more adaptable BTSP learning rates allowing neurons to form more robust place fields even at high speeds. Certainly, our findings suggest that place fields initially formed under outlier conditions, like very high or low running speeds, are likely to require additional BTSP learning events in order to fully stabilise, in contrast to place fields initially formed under more moderate conditions.

### 2.4 Place fields change shape if the target object is moved

We next tested how our model responded to the target landmark object being moved in the environment. Previous experiments have shown that when a second BTSP event is induced at a new location in a neuron that already has a place field, a new field forms at the new location. In some cases, the original place field is also dampened, in particular if the two locations are close to one another [Milstein et al., 2021]. To test this in our model, we ran simulations at our original running velocity of 0.25 m/s in which, after the initial BTSP induction event, we moved the landmark object to locations spanning the entire track (see Supp. Video 1). Once any additional BTSP weight updates had been applied, we ran evaluation traversals, and estimated average place fields as previously. It should be noted that moving an object can sometimes lead to broad remapping throughout the hippocampus, as if an animal had entered a new environment [Burke et al., 2011]. Here, our aim was to model cases in which the environment is rich enough that moving a single object does not lead to extensive remapping of the input place fields.

As expected, we found, in most cases, that moving the landmark object resulted in a BTSP-triggering plateau potential near the new location of the target. For distant locations, the resulting BTSP weight updates formed a new place field at the landmark object’s new location, with the original place field only decreasing slightly in amplitude. In contrast, for nearby locations, the original place field was more strongly dampened, or incorporated into an expanded place field encompassing both the previous and new target object location. In all cases, the peak of the place field shifted toward the landmark object’s new location. Notably, however, if the landmark object was moved very close to the original location (less than 0.2 m back or 0.3 m forward), no new BTSP events occurred, as the original place field was broad enough to block BTSP-triggering plateau potentials at the target’s new location (see Fig. 4 and S4). None of the simulations resulted in more than one additional BTSP event, indicating that the updated place fields were as stable as the original place field.

**Figure 4:**
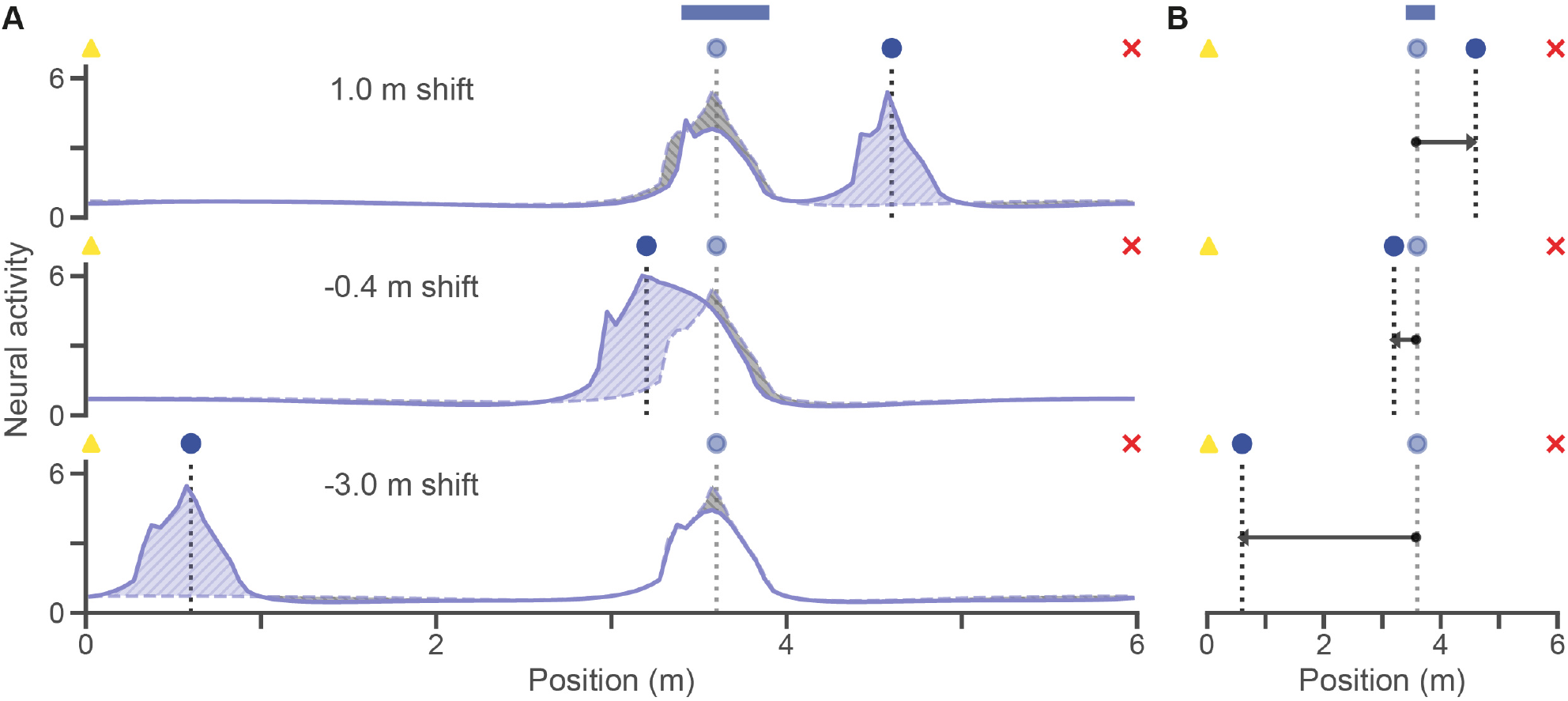
Moving the landmark object leads to additional BTSP events, changing place field shapes. **A**. Examples of place field changes after the landmark object was moved: 1.0 m shift (*top*), -0.4 m shift (*middle*), and -3.0 m shift (*bottom*). The original place field (dashed line) and the new place field (full line) are shown. Light purple shading and upward hatching (//) show increases in place field amplitude, whereas grey shading and downward hatching (*\\*) show decreases in place field amplitude. Icons at the top of each subplot show the start (yellow triangle) and reset (red x) positions. Original landmark object position (light blue circle and light vertical dashed line) and new position (blue circle and vertical dashed line) are shown for each example. Blue shading above shows the range of shifted positions that did not produce a second BTSP event (3.4 to 3.9 m, i.e., -0.2 to 0.3 m from the original landmark object position). **B**. Changes in place field peak location for the example place fields shown in **A**. Icons, dashed lines and blue shading at the top as in **A**. See Fig. S4 for results across the full range of object shifts.

Thus, our model successfully recapitulates experimental findings showing how consecutive BTSP events shape place fields, and makes specific predictions for how place fields that encode specific features of an environment should evolve if these features move. It also demonstrates how the hippocampus can adapt its predictive map through BTSP when features of an environment change.

### 2.5 Stable place field formation through BTSP extends to a 2D open field environment

Having established our model’s ability to recapitulate properties of place fields formed through BTSP in linear track simulations, we next tested whether the model’s self-stabilising place field formation could generalise to a 2D environment. Although BTSP has, to date, been studied primarily in 1D environments [Bittner et al., 2015, 2017; Dong et al., 2021; Milstein et al., 2021; Grienberger & Magee, 2022; O’Hare et al., 2022; Priestley et al., 2022; Rolotti et al., 2022; Xiao et al., 2023; Gonzalez et al., 2024; O’Hare et al., 2025; Neubrandt et al., 2025; Qian et al., 2025], the properties of place fields formed in the same circuits have been widely studied in 2D environments [O’Keefe & Dostrovsky, 1971; Mizuseki et al., 2012; Ziv et al., 2013; Moser et al., 2015]. Furthermore, 2D environments allow for more complex and diverse behaviours and navigation patterns, and are a better model of daily navigation of the world by humans and non-flying animals. It is therefore critical to confirm that our circuit can form stable predictive maps during navigation of a 2D environments.

For our first set of 2D simulations, we used a square open field `RatInABox` environment with one target landmark object. To retain some similarity to the linear track simulations, we first studied how the network would stabilise if the landmark object was consistently approached from the same general direction. To do this, we placed the object in a corridor at the bottom left of the open field, such that it was primarily reachable from the right (see Fig. 5A). The agent was initialised at a random location, most likely to be outside the corridor, and set to run at around 0.25 m/s (± 0.13 m/s). We guided the agent’s trajectory toward the target landmark object with some noise. To ensure good spatial coverage of the open field for estimating place fields, we included random walks between target reaches (see Fig. 5B). As in the linear track simulations, one object cell was initialized with a narrow field around the landmark object, providing a target to the pyramidal neuron. Place cell fields formed a grid across the entire environment (see Fig. 5C-D). Notably, only a few changes were made to the circuit parameters used in the linear track to compensate for the much larger number of input place cells (see Tables 1 and 2).

**Figure 5:**
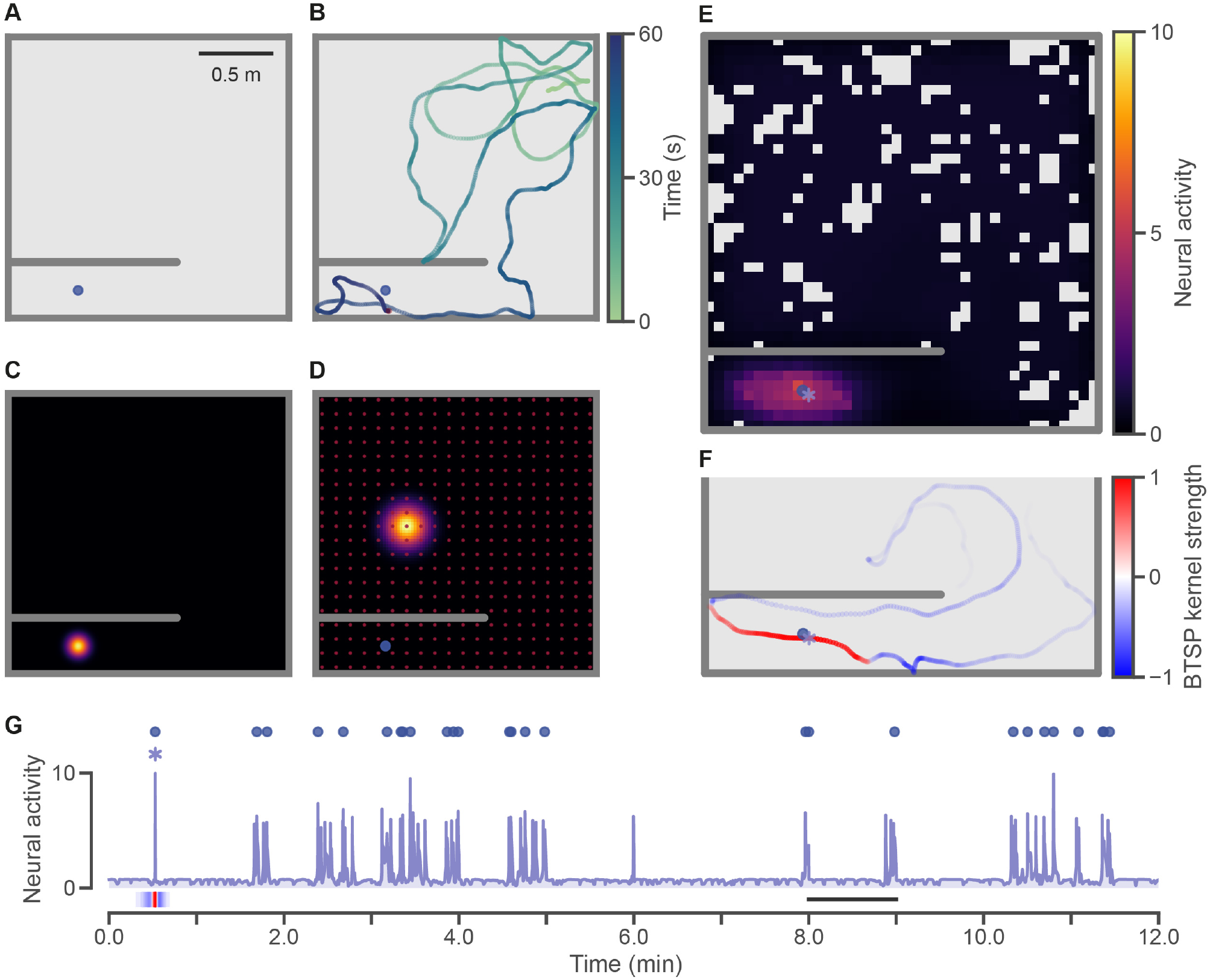
Place fields formed through BTSP are also self-stabilising in a 2D open field environment. **A**. 2D open field environment (2 m × 2 m). The landmark object is placed in a corridor in the bottom left, and primarily accessible from the right. Scale bar shows 0.5 m. **B**. Sample trajectory showing the agent’s path over one simulated minute. The trajectory goes from light green to dark purple as time advances. The agent’s final position is shown by a small red dot marking the end of the trajectory. **C**. Object cell field, centered on the landmark object (not shown). The object field is computed from neural activity expressed in arbitrary units ranging from 0 to 10. The colormap range is the same as in **E**. **D**. Place cell field centers are shown in red forming a 20×20 grid. One example place cell field is plotted around its place field center. The place field is computed from neural activity expressed in arbitrary units ranging from 0 to 10. The colormap range is the same as in **E**. **E**. Pyramidal neuron place field formed during an open field simulation. The place field is computed from neural activity expressed in arbitrary units ranging from 0 to 10. The light purple asterisk shows where the underlying BTSP event occurred. Spatial bins in the open field that were not visited by the agent during the time period used to compute the place field are plotted in light grey. **F**. Agent trajectory before and after the BTSP event (marked by a light purple asterisk) plotted as a function of the BTSP kernel’s strength at each time point. Trajectory dots are more transparent for kernel values near zero. For visual clarity, the kernel strength is plotted from -1 (most negative, blue) to 1 (most positive, red), but it should be noted that the negative and positive ranges are of different amplitudes (see Fig. 1D). **G**. Neural activity of the pyramidal neuron over the 12-minute simulation, expressed in arbitrary units ranging from 0 to 10. The time axis is in simulated minutes. A light purple asterisk shows when the BTSP event occurred. Each visit to the landmark object is shown by a blue circle. The BTSP kernel strength, shown in **F**, is plotted here as a function of time. The black line (from minute 8 to 9) marks the neural activity timecourse corresponding to the trajectory plotted in **B**.

In a 12-minute open field simulation, the pyramidal neuron formed a stable place field after experiencing one BTSP event during its first visit to the landmark object (see Fig. 5E). The place field shape reflected the agent’s trajectory before and after the BTSP event (see Fig. 5F and Fig. S5A-B), but was still general enough to prevent additional BTSP events from being triggered during the numerous additional visits to the landmark object that occurred throughout the rest of the 12-minute simulation (see Fig. 5G). To confirm the stability of few-shot place field formation with our model in a 2D environment, we ran nine additional open field simulations with different seeds. In four of these simulations, a second BTSP event was required for a stable place field to form (see Fig. S5C). In these cases, and in most subsequent simulations where BTSP was triggered more than once in nearby locations, weight normalisation was recruited to prevent runaway weight updates. Importantly, the place field formed in each simulation had a slightly different shape, reflecting the agent’s specific trajectory before and after each BTSP event occurred (see Fig. S5D). Overall, BTSP as implemented in our circuit model enables predictive, self-stabilising place fields to form in a 2D open field environment.

### 2.6 Place fields are reshapped by BTSP following changes to the transition structure of the environment

As with the linear track experiment, having shown that our circuit forms stable place fields in a 2D environment, we next tested how it would respond when the structure of the environment changed, after an initially stable place field had formed. If our model can indeed do predictive learning, it should be able to stably update its representations when their predictive power decreases due to changes to the environment. This time, instead of moving the landmark object, we changed the transition structure of the environment by introducing teleportation. Teleportation-like mechanisms are widely used in virtual reality experiments with rodents. For example, in linear track experiments, once an animal has reached the end of a virtual track, instead of having to retrace their steps, they will usually be “teleported” back to the start of the track or to a different track [Fiser et al., 2016; Gauthier & Tank, 2018; O’Hare et al., 2025]. In 2D virtual environments, teleportation ports could be used to improve our understanding of how spatial features are encoded in the hippocampus as they allow the behaviourally-relevant distance between features to be studied independently of their Euclidean, or straight-line, distance. For example, here, by introducing a pair of teleportation ports, we created a new way for the agent to approach the landmark object. We placed the entrance port in the upper open area of the open field, and made it left-facing, such that the agent had to come within range of it from the left in order to trigger a teleportation event. The exit port was positioned left of the landmark object, and was right-facing such that the agent exited it going right. When teleporting, the agent’s entry vector was reused as its exit vector, such that its exit locations were quite variable across teleportation events.

With this teleportation feature enabled, the agent could now rapidly reach the landmark object without approaching it gradually from the right, as it had previously done. The previous simulation (see Fig. 5) was extended until at least six teleportation events had occurred, and at least ten simulated minutes had elapsed since the last BTSP weight update was applied. The initial place field, which had been stable across repeated visits to the landmark object (see Fig. 5), was not able to prevent a BTSP event from being triggered following the agent’s first teleportation event, and was therefore updated to incorporate the area left of the teleport entrance port (see Fig. 6). The updated place field thus took into account the fact that the area near the teleport entrance port was behaviourally close to the landmark object, even though it was quite far in Euclidean distance. However, this first update was still not sufficient to account for the agent’s trajectory on the second teleportation event, which triggered another BTSP event (see Fig. 6). This final updated place field which displayed a stronger peak near the teleportation port entrance successfully prevented additional BTSP events from being triggered during the rest of the simulation. In particular, none of the four additional teleportation events recorded triggered additional plasticity, as, in each case, distal apical dendrite inhibition was able to ramp up sufficiently early to prevent additional plateau events from occurring (see Fig. 6 for the first four teleportation events, and see simulation #1 in Fig. S6 and Supp. Video 2 for the full simulation).

**Figure 6:**
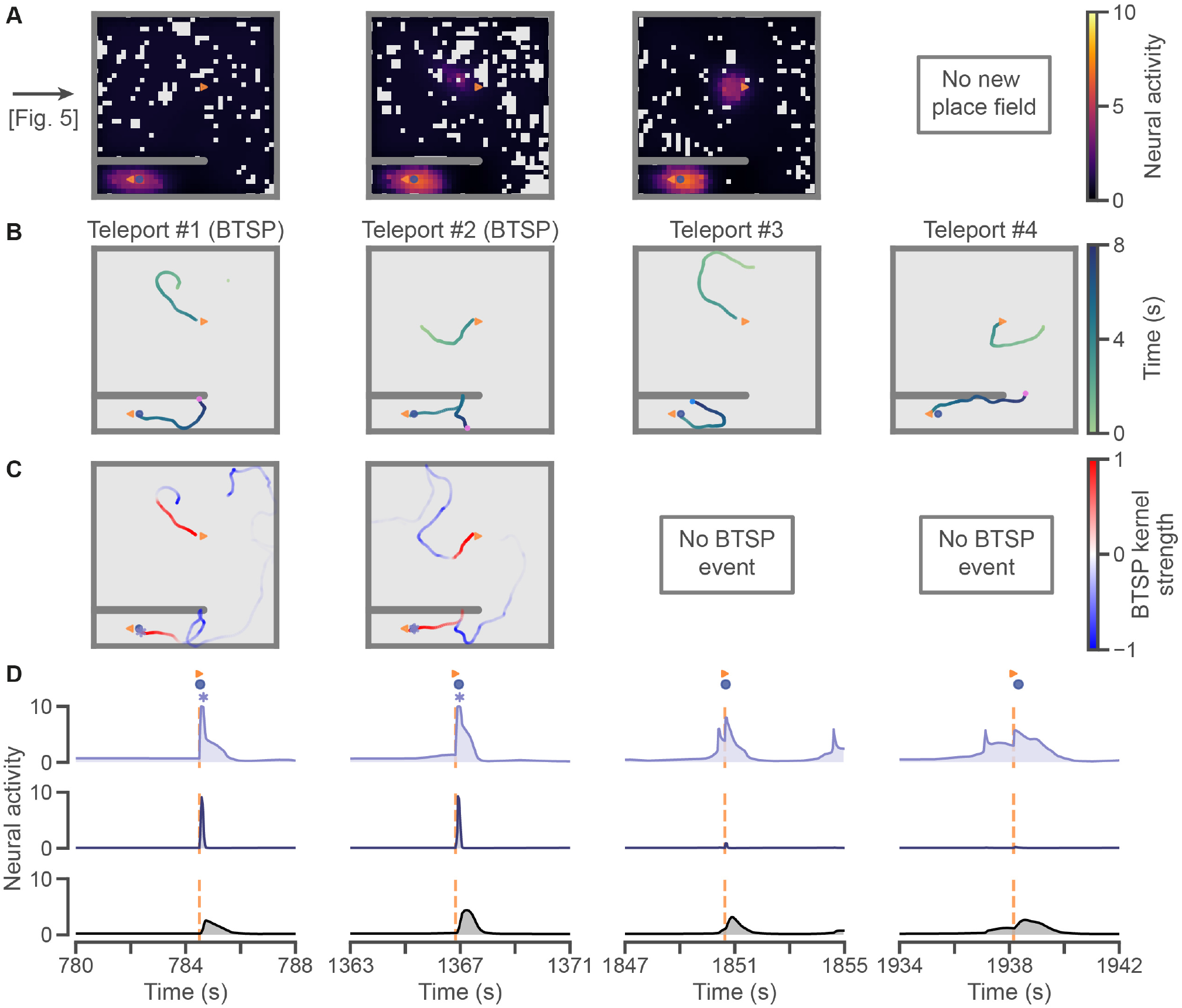
Place fields are updated through BTSP to reflect changes in the environment’s transition structure. **A**. Pyramidal neuron place field before each of the first four teleportation events. The first column shows the initial place field, before the first teleportation event, which is the same as shown in Fig. 5E. The second and third columns show the pyramidal neuron’s updated place fields following the BTSP events triggered by the first and second teleportation events. No new place field is plotted in the fourth column as the third teleportation event did not trigger additional plasticity. The fifth and sixth teleportation events recorded in the simulation, neither of which triggered additional BTSP events, are omitted for simplicity (see simulation #1 in Fig. S6). Place fields are computed from neural activity expressed in arbitrary units ranging from 0 to 10. The blue circle shows the landmark object, and the orange triangles show the teleportation ports. The left-facing entrance port is in the open area, and the right-facing exit port is next to the landmark object. **B**. Agent trajectory for eight simulated seconds around each of the first four teleportation events. The trajectory goes from light green to dark purple as time advances. The agent’s final position plotted is shown by a small pink dot. If a teleportation event triggered a BTSP event, this is noted in the trajectory plot title with “(BTSP)”. **C**. Agent trajectory before and after each BTSP event (shown by a light purple asterisk) plotted as a function of the BTSP kernel’s strength at each time point. Trajectory dots are more transparent for kernel values near zero. For visual clarity, the kernel strength is plotted from -1 (most negative, blue) to 1 (most positive, red), but it should be noted that the negative and positive ranges are of different amplitudes (see Fig. 1D). **D**. Neural activity for 8 simulated seconds around each of the first four teleportation events (as in **B**) for all three components of the pyramidal neuron circuit: the proximal compartment (light purple, *top*), the distal compartment (dark purple, *middle*) and the inhibitory interneuron (black, *bottom*). Light purple asterisks mark BTSP events. Teleportation events are shown with a vertical orange dashed line aligned to an orange triangle above the proximal compartment activity. Blue circles mark landmark visits.

To confirm the reliability of this phenomenon where place fields are stably updated to reflect changes in the transition structure of the environment, we ran seven additional simulations with different seeds. In these simulations, we also varied the exact position and orientation of the teleportation ports (see Fig. S6). In all but two simulations (#5 and #8), the initial place field self-stabilised after one BTSP event. Once teleportation was enabled, additional BTSP events were triggered in all simulations, almost always following a teleportation event. In most of the simulations, only one or two additional BTSP events were required for the updated place field to stabilise. In three simulations, however, the pyramidal neuron underwent three to four additional BTSP events before forming a stable place field that could fully account for the wider variety of ways in which the agent could now reach the landmark object. Nonetheless, the number of BTSP events was in all cases significantly lower than the number of landmark visits and lower than the total number of teleportation events (see Fig. S6). Importantly, all of the final place fields reflected the new structural feature of the environment whereby the area from which the agent could enter the teleportation entrance port was much more predictive of encountering the landmark object than the rest of the upper open field area. Together, these findings show how BTSP, as implemented in our network, allows a place field to be stably updated to reflect changes in the transition structure of the environment, thus ensuring it continues to be sufficiently predictive of the feature it anticipates. The broad range in results observed also further confirms the close relationship in our model between an agent’s spatial behaviour and the properties of the place fields it learns (see Fig. S6). As in the linear track experiments, our model predicts that it is the agent’s early or unpredictable encounters with a feature that most strongly shape the place fields its pyramidal neurons learns. Since later encounters of a feature are increasingly likely to be well-predicted by previous encounters, our model supports experiments showing that they play a reduced role in shaping the properties of the place fields learned in an environment [Pecirno & Keinath, 2025].

### 2.7 Place field formation is stable in a population of neurons with distinct targets

An important feature of our simulations was that although they were necessarily discrete in time, our agent’s trajectories were designed to be realistic, variable in speed and direction, and continuous in space [George et al., 2024]. This greatly increased the number of ways in which a target object could be approached by an agent, in particular in comparison to gridworld simulations, and thus made the task of learning predictive, self-stabilising place fields more challenging. To confirm that our circuit could successfully and robustly predict environmental features under these more complex, but realistic conditions, we tested it in a multi-target open field task. For this simulation, we initialised an open field with four internal walls and 40 landmark objects scattered across the whole area. Accordingly, we expanded our pyramidal neuron population to comprise 40 neurons (one per landmark object), each with its own inhibitory neuron and a different object cell targeting its distal compartment. To further demonstrate the robustness of our model, we added noise to the initial input place cell weights and to the proximal compartment’s neural activity. We then simulated the agent running from landmark object to landmark object for just over two simulated hours (see Fig. 7A). All 60 BTSP events recorded occurred within the first 110 minutes and neural activity from the final 15 minutes was used to compute place fields for each pyramidal neuron. By the end of the simulation, place fields covered most of the open field (see Fig. 7B-C and Supp. Video 3), with all 40 pyramidal neurons forming a place field around their respective target objects (see Fig. 7D-E and S7). Although the total number of visits reached almost 150 for some landmark objects, almost all neurons only underwent one or two BTSP events before forming a stable place field. Three neurons required more than two BTSP events, but this was not linked to a higher number of visits (see Fig.7D and S7C). Thus, the results of this final multi-target simulation confirm our hippocampal circuit’s ability to produce stable 2D place fields using realistic trajectories through spatially continuous environment like the ones experienced under real experimental conditions. Finally, the ability of place fields learned from only a few BTSP events to account for hundreds of different visits to the same object confirms the strong predictive power conferred by the broad timecourse of the BTSP kernel and its suitability to drive predictive learning in the hippocampus.

**Figure 7:**
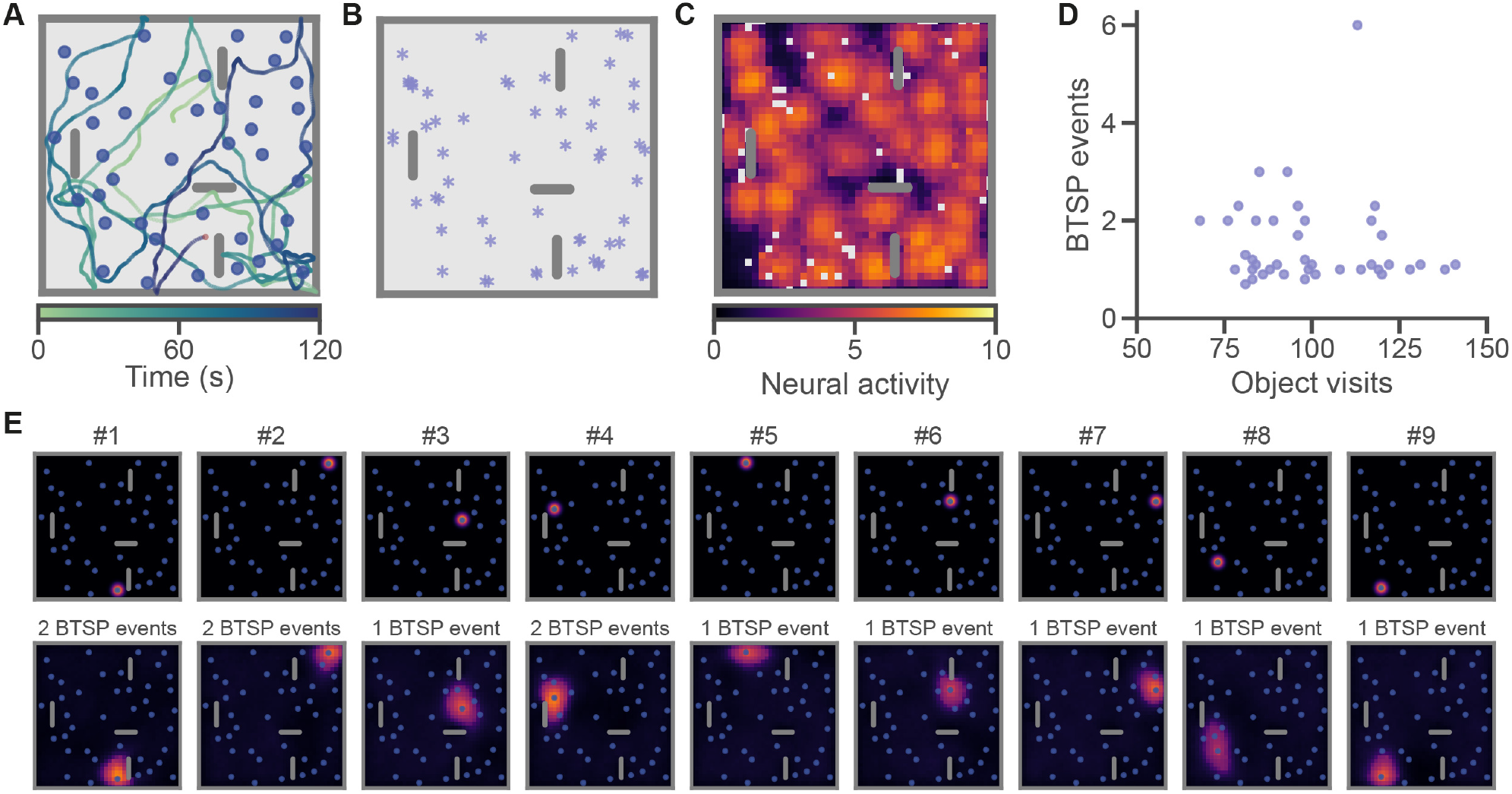
Place field formation through BTSP is stable across a population of pyramidal neurons with different target objects. **A**. Agent trajectory over the first two simulated minutes around an open field with 40 landmark objects and four internal walls. The trajectory goes from light green to dark purple as time advances. The agent’s final position is shown by a small red dot. **B**. Locations of all 60 BTSP events recorded in the pyramidal neuron population during the multi-target open field simulation. **C**. Pyramidal neuron place fields at the end of the simulation, overlayed for all 40 pyramidal neurons. Each place field is computed from neural activity expressed in arbitrary units ranging from 0 to 10, and the maximum value across all pyramidal neurons is plotted. Spatial bins in the open field that were not visited by the agent during the time period used to compute the overlayed place fields are plotted in light grey. **D**. Relationship between the number of target visits and the total number of BTSP events recorded for each pyramidal neuron. The number of BTSP events can only take integer values, but some jitter is added to the plotted values to ensure dots do not overlap. **E**. Target object cell field (*top*) and place field at the end of the simulation (*bottom*) for each of nine pyramidal neurons. The number of BTSP events recorded for each of these pyramidal neurons is recorded above the place field plot. The object and place fields are computed from neural activity expressed in arbitrary units ranging from 0 to 10. The colormap range is the same as in **C**. For place fields for all 40 pyramidal neurons, see Fig. S7A.

## 3. Discussion

Predictive learning is a leading hypothesis for how the brain forms cognitive maps of environments and guides behaviour. Research into this hypothesis has focused heavily on sensory processing areas, and the circuitry properties specific to these regions. Although the hippocampus shares many of these properties and is thought to play a central role in predictive learning, circuit-level implementations have not yet been thoroughly examined. Here, we demonstrated how predictive learning emerges directly from key properties of hippocampal circuits and enables stable behaviourally-relevant features to be learned in a few-shot manner. First, we demonstrated how stable place fields around target objects could be formed in a linear track simulation using a simple circuit centered on a self-inhibiting two-compartment CA1-like pyramidal neuron subject to internally-triggered BTSP-like plasticity. The place fields learned by these two-compartment pyramidal neurons showed key features observed experimentally in hippocampal research. Place fields were formed in a one or few-shot manner [Bittner et al., 2015, 2017; Milstein et al., 2021; O’Hare et al., 2022; Gonzalez et al., 2024; O’Hare et al., 2025; Madar et al., 2025b; Dorian et al., 2026]. Place field width was correlated to running velocity during place field formation [Bittner et al., 2017; Milstein et al., 2021; O’Hare et al., 2022; Rolotti et al., 2022; Gonzalez et al., 2024; O’Hare et al., 2025], and the effect of a second BTSP event on place field shape was distance-dependent [Milstein et al., 2021]. Secondly, we confirmed that the same circuit, with only slightly modified parameters (see Tables 1 and 2), also enabled one or few-shot formation of stable place fields in a two-dimensional continuous environment. By adding teleportation ports partway through learning, we showed that the circuit was able to update its place fields in response to changes in the environment’s spatial structure so as to better represent the predictive relationship between different spatial locations and the target landmark object. Lastly, we demonstrated that our circuit could support few-shot learning of a broad set of stable place fields for differently positioned and visited target landmark objects, as would be encountered during naturalistic exploration of a realistic environment.

### 3.1 Comparisons to the experimental literature and model predictions

Our model is consistent with key phenomena reported in the experimental literature, and makes several predictions for future experiments. First, our circuit posits a critical role in feature learning for hippocampal interneurons and particularly OLM interneurons which target distal apical dendrites [Geiller et al., 2020; Milstein et al., 2021; Udakis et al., 2025]. This is consistent with recent work showing that OLM interneurons play a key role in gating plateau potentials in CA1 pyramidal neurons [Vaasjo et al., 2025]. Increasing their activity not only reduces the frequency of plateau potentials in pyramidal neuron cell bodies, but also interferes with place field formation. In contrast, decreasing their activity late in learning increases new place field formation. These results are consistent with the effects observed in our circuit, where increasing the inhibitory weight prevented BTSP learning, and decreasing it prevented self-stabilisation of existing fields (see Fig. S2). Since the inhibitory interneurons in our model inherit place fields from their paired pyramidal neurons, our model is also consistent with the finding that OLM neurons show an increase in activity over learning [Sheffield et al., 2017; Arriaga & Han, 2019; Geiller et al., 2020; Hainmueller et al., 2024; Udakis et al., 2025, though see Kipper et al., 2025], develop place fields consistent with those measured in the pyramidal neuron population, and remap in new environments [Campbell et al., 2025; Kipper et al., 2025]. Our model additionally predicts that the output of OLM neurons should be delayed with respect to the output of the pyramidal neurons that drive their activity, forcing pyramidal neurons to develop fields that are meaningfully predictive of EC3 inputs to their distal apical dendrites. Our circuit model also supports a key role for EC3 neurons in shaping what is predictively encoded in CA1. For example, activating more EC3 inputs in response to specific sensory features would be expected to lead to overrepresentation in CA1. In real animals, this might bias behaviour toward these locations if overrepresentation plays a direct role in guiding behaviour. Consistent with this, studies have shown that inhibiting EC3 activity largely abolishes the enrichment of place fields around landmark locations that is typically observed during learning, and impairs learning in spatial and temporal delay tasks [Suh et al., 2011; Grienberger & Magee, 2022].

The most critical prediction our model makes is that changes to an environment should lead to error-like signals, reflecting a significant increase in BTSP-triggering plateau potentials, specifically around features whose predictability has changed. In this vein, our simulations are already consistent with the finding that cognitive maps in CA1 are most strongly reflective of early learning experiences rather than later ones [Pecirno & Keinath, 2025]. In our model, this is because early experiences in an environment are generally less well predicted by virtue of being novel, and thus drive significant BTSP-driven learning. However, both our object shift and teleportation simulations also provide clear examples of how later changes to an environment, if they do not trigger complete remapping, should also lead to an increase in BTSP-driven learning. For example, in our simulations, the addition of a teleportation port critically changes how the agent is able to reach the landmark object, and thus leads to additional BTSP-triggering plateau potentials around the exit port. As a result, place fields situated near the exit port are updated to incorporate locations near the entrance port. Although numerous computational studies have proposed a role for the hippocampus in predictive learning [Lisman & Redish, 2009; Stachenfeld et al., 2017; Santos-Pata et al., 2021; Chen et al., 2024; Levenstein et al., 2024; Bennett et al., 2025; Liu et al., 2025], comparatively few studies have been conducted to look specifically for predictive learning signals in hippocampal neurons. Several studies have shown evidence of anticipatory signals in the hippocampus [Muller & Kubie, 1989; Mehta et al., 1997; Ferbinteanu & Shapiro, 2003; Lee et al., 2006; Pastalkova et al., 2008; Lisman & Redish, 2009], but much of the evidence for error-like signals comes from human studies in which, in most cases, responses cannot be characterised at the level of individual neurons [Aitken & Kok, 2022; Pecirno & Keinath, 2025; Bein et al., 2020; Kok et al., 2020; Kumaran & Maguire, 2006, though see Aquino et al., 2024]. Lee et al. [2017] and Yaghoubi et al. [2026] do report reward prediction-like error signals in the hippocampus. However, the focus of these studies is on value learning in the context of a reward task, and not on learning to predict features in a changing environment. Studies specifically investigating the effects of changing the relational structure of a task are critically needed to confirm the hippocampus’ role in predictive learning and to validate models like ours. We predict that such experiments would show an increase in BTSP-triggering plateau potential frequency specifically in neurons whose activity is predictive of the affected features, as well as changes in these neuron’s fields to incorporate the new information.

### 3.2 Limitations and opportunities for future development

While our model captures several important features of the hippocampal circuitry, with certain exceptions discussed below, there are certain limitations to its biological plausibility. First, the one-to-one mapping of OLM-like inhibitory interneurons to pyramidal neurons is not biologically plausible. Simplifications like these were made to create a model that is interpretable and provides a proof of concept for feature learning in the hippocampus. However, in reality, the circuitry is of course much more complex. Single OLM inhibitory interneurons receive input from numerous pyramidal neurons, and target the distal apical dendrites of numerous pyramidal neurons in return [Topolnik & Tamboli, 2022]. Expansions to our circuit model could test the effect of weakening the one-to-one constraint. In this case, the one-to-one mapping of EC3 inputs to pyramidal neurons would likely also need to be weakened or removed, and competitive mechanisms like lateral inhibition might be needed to ensure that different pyramidal neurons form different fields [Haga & Fukai, 2018; Rolotti et al., 2022]. This would likely reduce the circuit’s interpretability, but certainly improve its plausibility. In the long-term, future models will hopefully also determine how complexly organized inputs from EC3 interact with OLM inhibition at the level of individual dendritic branches to generate the localized responses that, when properly coordinated, lead to large dendritic events like plateau potentials [Moore et al., 2021b; O’Hare et al., 2025]. Models like these could provide key insights into how CA1 pyramidal neurons learn to encode highly conjunctive features of the environment.

Although our implementation of BTSP-driven weight updates is derived from experimental data, it does differ in certain ways from what was proposed in seminal work measuring and modeling BTSP [Milstein et al., 2021]. In this work, BTSP was implemented for a spike-based model as a weight-dependent rule scaled by two time-dependent factors: the product of an exponentially filtered eligibility trace corresponding to presynaptic spikes, and an instructive signal corresponding to postsynaptic dendritic plateau potentials. In our rate-based model, plateau potentials were also used as discrete instructive signals. However, since our model was not a spiking model, we used the presynaptic neural activity rates to compute the eligibility traces. Our BTSP kernel was then applied by filtering the inputs to each pyramidal neuron using differences-of-exponentials fit to the BTSP kernel timecourse reported by Milstein et al. [2021] (see Fig. 1D). Each filtered input was then integrated and multiplied by the BTSP update factor to yield the value by which the input’s weight onto the pyramidal neuron would be incremented. Notably, this makes our learning rule technically activity-dependent instead of being weight-dependent. Since, in our simulations, pyramidal neural activity was heavily driven and thus closely related to input place field weight, this resulted in a very similar learning rule. However, to limit synaptic potentiation and depression, and drive updates toward stable target weights, as has been observed experimentally, we added weight normalisation. Specifically, we applied divisive normalisation when the weights of a pyramidal neuron exceeded a specific threshold [Mainali et al., 2025]. We believe that our approach presents a good compromise as it produces BTSP updates that greatly resemble those reported experimentally, while also being easily applied online using rate-based neurons. Nonetheless, modifying the learning rule to more closely match the one proposed by Milstein et al. [2021] might improve the model’s ability to capture experimentally-reported phenomena like the relationship between running velocity and place field width which, at higher speeds, is not fully reflected in our model (see Fig. 3).

Several features of the hippocampal circuit were also omitted in our model. For example, we did not model other inhibitory interneuron subtypes like parvalbumin interneurons, vasoactive intestinal peptide interneurons and bistratified somatostatin interneurons which certainly contribute to hippocampal computations [Arriaga & Han, 2019; Geiller et al., 2020; Hainmueller et al., 2024; Kipper et al., 2025; Neubrandt et al., 2025] whose analogs are thought to play key roles in predictive learning in sensory cortex [Garrett et al., 2020; Hertäg & Sprekeler, 2020; Aizenbud et al., 2026]. We also did not include more traditional, shorter timecourse plasticity like Hebbian learning [Bi & Poo, 1998]. If incorporated into our model, Hebbian learning would likely allow the circuit to slowly reshape BTSP-induced place fields across learning. This could enhance their predictive power by ensuring that a place field’s shape reflects not only the agent’s first visit to a landmark object, but also the broader statistics of its subsequent visits [Bono et al., 2023; George et al., 2023b]. In agreement with this, Madar et al. [2025a] report that whereas BTSP best explains the formation of new place fields, Hebbian learning best explains how place field properties shift across learning. Lastly, as mentioned above, our simple two-compartment model cannot fully capture the complex electrical properties of a CA1 pyramidal neuron with its complex branching apical dendritic tuft, radial obliques and basal dendrites. More biophysically realistic models are likely needed to explain phenomena observed locally in dendritic branches, like the increase in plateau potentials measured in tuft dendrites following place field formation reported by O’Hare et al. [2025] and to determine how these relate to the BTSP-triggering plateau potentials typically measured at the cell body which our model focuses on.

The role of neuromodulation and oscillations, in the theta band for example, is also not considered in our model. Both are thought to play a key role in enabling the hippocampus to serve its two critical, but in some ways opposite roles: the encoding of new memories and the retrieval of stored memories. Notably, both theta oscillations and the release of neurotransmitters likely acetylcholine shape the relative influence of EC3 and CA3 inputs on pyramidal neuron activity in CA1. For example, EC3 and CA3 inputs tend to be prioritised at different phases of the theta cycle [Brankačk et al., 1993; Ang et al., 2005; Hasselmo, 2006; Fernández-Ruiz et al., 2017; George et al., 2023a]. Relatedly, acetylcholine, which is involved in shaping theta oscillations, specifically increases the priority of EC3 over CA3 inputs [Palacios-Filardo et al., 2021], and is delivered to CA1 in response to negative and positive rewards [Lovett-Barron et al., 2014; Teles-Grilo Ruivo et al., 2017] as well as when novel environments are first being explored [Xuan et al., 2025]. Together, mechanisms like these may allow the hippocampus to shift toward memory reactivation, prioritising CA3 inputs when there is sufficient evidence that an environment is familiar, and toward memory encoding, prioritising EC3 inputs, otherwise [Cabral et al., 2014; Kaifosh & Losonczy, 2016; Wang et al., 2025]. In our model, we considered only memory encoding and did not investigate the hippocampal circuitry’s role in memory retrieval. We expect, however, that this role heavily involves CA3. For simplicity, we modeled CA3 as providing a purely spatial backbone based on which predictive features are learned in CA1. This is because predictive features have been primarily reported in CA1 [Muller & Kubie, 1989; Mehta et al., 1997; Ferbinteanu & Shapiro, 2003; Lee et al., 2006; Pastalkova et al., 2008], and although CA3 also undergoes BTSP learning, its learning kernel does not share the asymmetrical shape observed in CA1 [Milstein et al., 2021]. However, with its highly recurrent circuitry, CA3 is particularly well-structured to support pattern completion. This points to a critical role in memory retrieval and in sequence learning through replay, both of which are likely required for the formation of coherent cognitive maps [O’Reilly & McClelland, 1994; Neunuebel & Knierim, 2014; Guzman et al., 2016; Mishra et al., 2016; Levenstein et al., 2024]. Future work could expand the role of CA3 and enable the circuit to switch dynamically between memory formation and reactivation. Such work might also benefit from incorporating the subiculum which is the primary output of the hippocampus and has been reported to encode predictive features with longer time horizons than CA1 [Bennett et al., 2025].

There are also certain phenomena observed in the hippocampus that we did not model here. One such phenomenon is the disappearance of place fields during exploration [Zheng et al., 2024]. The BTSP kernel we applied accounts for one-shot formation of place fields, but not for one-shot disappearance. As seen in Fig. 4, after an initial place field is formed, subsequent BTSP updates can partially depress the original place field. However, it would likely take multiple BTSP events to fully abolish a place field using our BTSP kernel. Milstein et al. [2021] do report examples where induction of a second BTSP event appears to fully abolish the original place field. A probabilistic implementation of BTSP where only the kernel’s expected shape matches the parameters reported by Milstein et al. [2021], and the exact shape at individual induction events varies might be required to suitably capture this type of phenomenon.

Importantly, our simulations, and the majority of our discussion, have focused on learning spatial features in CA1, even though the hippocampus forms cognitive maps that encode a variety of sensory, reward-related and even temporal features, as well as complex conjunctions of these features [Terada et al., 2022]. This focus was chosen in part because spatial tasks are among the most widely studied in the hippocampus. In addition, the fields encoded by pyramidal neurons trained on continuous tasks, like spatial learning, lend themselves particularly well to visualisation, comparison and fine-grained analysis. Nonetheless, in future work, our circuit could be applied to study learning of predictive, but non-spatial features. For example, a task could be tested in which certain targets encoded by entorhinal inputs to the distal compartments are preceded by a specific olfactory or visual cue [Zhao et al., 2022; Dorian et al., 2026]. As long as CA1 neurons receive inputs to their proximal compartments signaling these cues, they should be able to learn to anticipate the targets using features that are not strictly spatial. Conjunctive tasks could also be assessed in which, for example, a target is predicted by an olfactory cue presented in a specific spatial location. Such experiments could also shed further light on the relationship between hippocampal and sensory predictive learning. In sensory cortices, predictive learning is often studied over very short time dependencies, even on the order of milliseconds. In contrast, the hippocampus is thought to learn more abstract, behaviourally-dependent predictive relationships unfolding over seconds or more. Thus, it is possible that short-term sensory predictions in early sensory areas are guided by much more complex and broad predictions which originate in high level areas like the hippocampus. These might then be transformed into predictions that are more temporally and spatially localised, but also more sensorially specific as they are transmitted down the sensory hierarchy [Kumaran & Maguire, 2006; Keller & Mrsic-Flogel, 2018; Schlossmacher et al., 2022; Aizenbud et al., 2026]. Multi-area experiments on predictive learning are needed to evaluate whether this is indeed the case and how the hippocampus is involved.

Finally, and related to the discussion above on memory reactivation, we did not attempt here to address the question of how multiple maps can be learned in the hippocampus, without sequentially erasing one another [Sheintuch et al., 2020; Remme et al., 2021; Fenton, 2024]. One promising hypothesis is that the hippocampus uses different neuron ensembles or even synapse ensembles within the same neurons for different environments [Kubie & Muller, 1991; Treves & Rolls, 1994; Fenton, 2024; O’Hare et al., 2025]. For example, in each new environment, rapid learning, under the control of local inhibition, neuromodulation and intrinsic excitability, might be enabled for only a subset of CA1 neurons and a subset of the radial oblique dendrites targeted by CA3 neurons. When an animal moves into a new context and the hippocampus undergoes remapping, the subset of synapses or neurons subject to BTSP may be changed, temporarily protecting recently acquired cognitive maps from being overwritten. During this time, these newer cognitive maps stored in the radial oblique synapses may be transferred to more slowly changing synapses [Remme et al., 2021], like the basal dendrite synapses of CA1 neurons. These dendrites are also innervated by CA3, but their synapses appear to undergo more gradual changes than the synapses onto radial oblique dendrites [Gonzalez et al., 2024]. This could explain how cognitive maps stored in the hippocampus avoid being overwritten in the medium term before they are incorporated through consolidation to long-term memory in the prefrontal cortex [McClelland et al., 1995; O’Reilly et al., 2011; Srinivasan et al., 2023]. Future experiments exploring these ideas could modify the CA1-like pyramidal neurons by splitting the single proximal compartment into multiple ones for parallel storage of cognitive maps and a basal compartment for medium-term storage. The circuit’s ability to create and maintain several cognitive maps in parallel could then be assessed.

In summary, our model provides a compelling account of how key properties of the hippocampal circuit, namely BTSP, the complementary streams that target the distal and more proximal dendrites, and distal dendrite targeted inhibition, can together explain the hippocampus’ proposed central role in predictive learning. Our work also sets forth several key predictions whose experimental verification is needed to validate this increasingly prominent hypothesis. The codebase used for all simulations reported here is openly available for reuse and was built with the `RatInABox` toolbox which allows users to simulate continuous one or two-dimensional spatial trajectories for rodent-like agents along with the activity of a wide range of spatially-modulated neurons [George et al., 2024]. Thus, our work also offers an excellent starting point for researchers looking to delve further into the role of these key properties of the hippocampal circuit in the learning and memory-related phenomena studied here, and beyond.

## 4 Methods

### 4.1 Hippocampal circuit model

We simulated neural activity in a circuit model of the rodent hippocampus using `RatInABox`, an open-source Python toolkit for modeling realistic rodent locomotion and generating synthetic rate-based neural data for spatially modulated neurons [George et al., 2024]. The circuit comprised three areas: (1) a place cell area, (2) an object cell area, and (3) a pyramidal neuron area. In all areas, we modeled neurons as rate units with neural activity levels *R* ranging from 0 to 10 (arbitrary units). We modeled all neurons, except pyramidal neurons described below, as single compartment neurons.

#### Main circuit

We implemented the neurons in the place cell area and in the object cell area as`RatInABox PlaceCells` [George et al., 2024]. Place cell centers for the *N*_P_ neurons in the place cell area tiled each environment uniformly (60 in the linear track environment, 400 in a 20×20 grid in the open field environment). Each place cell’s activity was determined by the agent’s position with respect to a Gaussian distribution (st. dev. 0.1 m) centered on its place cell center [Mizuseki et al., 2012; Ziv et al., 2013]. The object cell area comprised one neuron for each simulated object and its neural activity was determined by the agent’s position with respect to a narrow Gaussian field (st. dev. 0.05 m) centered on the object neuron’s target object [Grienberger & Magee, 2022]. For both place and object cells, we used “line-of-sight” geometry such that spatial fields were bounded by nearby internal walls, if present.

#### Proximal compartment

The pyramidal neuron area comprised one pyramidal neuron for each landmark object in the environment (*N*_O_), and a paired inhibitory interneuron. Pyramidal neurons were modeled as two-compartment neurons with a distal compartment coupled to a proximal compartment. The proximal compartment of each pyramidal neuron *i*, representing its cell body and radial oblique dendrites, received as input the neural activity *R*_P_ from all *N*_P_ neurons in the place cell area, modulated by plastic weights 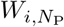 (from the weight matrix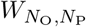). The weights were all initialised at 0.1 (linear track environment) or 0.04 (open field environment). These parameter values were chosen as they were just strong enough to enable initial learning to reliably occur around landmark objects. The proximal compartment also received as input the neural activity *R*_D_ of its paired distal compartment, with a fixed weight of 1.0. This ensured that strong neural activity in the distal compartment, which was modeled to be more bimodal (see below), was consistently propagated to the proximal compartment.

To enable plateau potential dynamics to emerge in the pyramidal neurons, we also gave each pyramidal neuron compartment an NMDA receptor-mediated current (*I*_S-NMDA,*i*_ for the proximal compartment), described in the next section. The proximal compartment’s neural activity *R*_S,*i*_ was then computed by summing the weighted inputs and passing the result through a sigmoid nonlinearity *σ*_S_. To approximate a linear function around its midpoint, we set the sigmoid’s width to 8.0 and midpoint to 6.0. We then projected its output into the 0 to 10 range. Neural activity for the proximal compartment was therefore computed as:

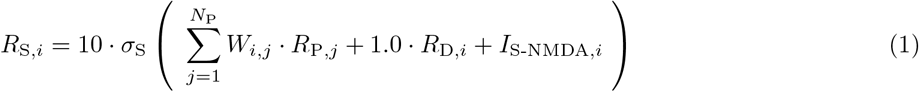

#### Distal compartment

The distal compartment of each pyramidal neuron *i* received as input theneural activity *R*_O,*i*_ of a paired neuron in the object cell area, with a fixed weight of 0.4, to drive strong activation near the landmark object. It also received as input the neural activity *R*_S,*i*_ of its paired proximal compartment with a fixed weight of 0.2. This weight was set at a lower value than the complementary distal to proximal weight based on experiments showing that action potential propagation into the distal apical tuft of CA1 neurons is attenuated at dendritic branch points and during spike trains [Spruston et al., 1995; O’Hare et al., 2025; Gonzalez et al., 2026]. Lastly, the distal compartment received as input the neural activity *R*_inh,*i*_ of its paired inhibitory interneuron with a fixed weight of 1.0. As with the proximal compartment, we gave the distal compartment an NMDA receptor-mediated current *I*_D-NMDA,*i*_. We then computed the distal compartment’s neural activity *R*_D,*i*_ by passing the summed weighted inputs through a sigmoid *σ*_D_. Here, we chose a narrower sigmoid (width: 6.0; midpoint: 5.0) to reflect the more bimodal activity propagation properties of dendrites compared to the cell body [Spruston et al., 1995], and projected the output into the 0 to 10 range.

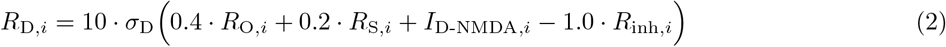

#### Inhibitory interneurons

To ensure that the hippocampal circuit learned to anticipate landmark objects, we introduced a time-delay in the distal inhibition. To do this, we computed an exponentially filtered version *F*_S,*i*_ of the neural activity of the proximal compartment at each time point *t* using a time constant of 0.3 sec (*α*_inh_ ≈ 0.095, filter initialised at 0), except in Fig. S1 where we assessed the effect of changing certain parameter values on place field metrics:

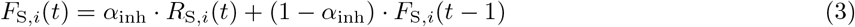

Each inhibitory interneuron *i* received as input the filtered neural activity *F*_S,*i*_ of the proximal compartment of its paired pyramidal neuron *i*, modulated by a fixed weight of 1.0 (except in Fig. S1, and in Fig. S2 where we visualised the effect of three values, 0.5, 1.0 and 2.0, on circuit activity). Finally, to compute each inhibitory interneuron’s neural activity, we passed the result through the same sigmoid *σ*_S_ as we used for the pyramidal neuron’s proximal compartment, and projected the output into the 0 to 10 range:

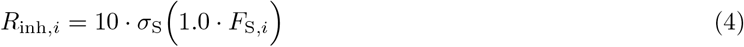

#### Simulation noise

In the multi-target open field task, noise was added to the simulation in two forms. First, instead of initialising all of the weights from the place cell neurons to the proximal compartments 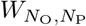 to 0.04, the weights were sampled from a Gaussian distribution with mean 0.04 and standard deviation 0.1. Additionally, background noise sampled from an Ornstein Uhlenbeck process, i.e., a stationary Gaussian Markov process, was added to the neural activity of the proximal compartment at each step by setting the neural layer’s `noise scale` parameter to 0.1 instead of 0 as in the other simulations [George et al., 2024].

#### Neural activity updates

We ran all simulations at around 33 Hz, i.e., with neural activity updated in steps with a Δ*t* of 0.03 seconds. At each time step, the agent’s position was updated as described below. Neural activity was then updated in this order: (1) place cells, (2) object cells, (3) inhibitory neurons, (4) NMDA receptor-mediated currents, (5) pyramidal neuron distal compartments, (6) pyramidal neuron proximal compartments. Circuit parameters are summarized in Tables 1 and 2.

### 4.2 NMDA receptor-mediated currents

To confer more complex internal dynamics to the pyramidal neurons and enable them to generate plateau potentials, we simulated NMDA receptor-mediated currents in both compartments. In real brains, NMDA receptors act as coincidence detectors, as they are both ligand and voltage-gated [Nowak et al., 1984]. Once activated, they remain active for an extended period of time, then become temporarily desensitised before they can be reactivated. Thus, NMDA receptors, although less involved in initiating neural activity, play a key role in amplifying and sustaining neural activity, critically shape neural activity dynamics [Lester et al., 1995] and are key drivers of plasticity, including BTSP [Tsien et al., 1996; Remy & Spruston, 2007; Grienberger et al., 2014; Bittner et al., 2017].

#### Dynamic variables

To simulate the NMDA receptor-mediated currents, we first simulated the evolution of three variables across time for each compartment: (1) the proportion *P*_BD_(*t*) of NMDA receptors bound at time *t*, (2) the proportion *P*_DS_(*t*) of NMDA receptors desensitised at time *t*, and (3) the proportion *P*_ACT_(*t*) of NMDA receptors activated at time *t*. At each time step, we first computed intermediate values 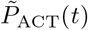 and 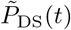 by decaying the values recorded at time *t* − 1 using exponential time constants of 0.1 sec and 0.3 sec, respectively (filters initialised at 0) [Lester et al., 1995]. Next, we computed *P*_DS_(*t*) by converting the decayed activation to desensitisation (the same equation applies to both compartments):

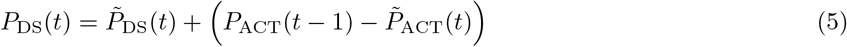

#### Bound NMDA receptors

We then updated *P*_BD_(*t*). Given that NMDA receptor binding is determined by the ligand concentration at the synapse, but not the synaptic weight, we modeled the binding proportion as a function of the sum of input activity (*R*_P_ or *R*_O_) targeting the specific compartment (proximal or distal, respectively) of neuron *i* at time *t*. To convert this value into a proportion, we applied a wide sigmoid function *σ*_BD_ (width set to 20 and midpoint at the two-thirds point, around 13.3):

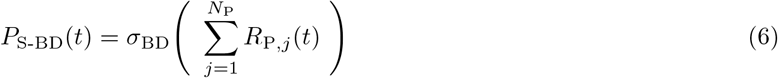

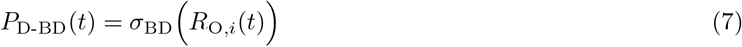

#### NMDA receptor activation

Lastly, we updated *P*_ACT_(*t*). As explained above, NMDA receptors are voltage-gated, in addition to being ligand-gated. Thus, for inactive, but bound NMDA receptors to activate, the compartment must pass a neural activity threshold. To compute the proportion of new NMDA receptor activation, we passed the current firing rate of each pyramidal neuron compartment *R*(*t*) (either *R*_S_(*t*) or *R*_D_(*t*)) through a narrow sigmoid *σ*_ACT_ (width: 0.5, distal midpoint: 0.8, proximal midpoint: 2.0). Midpoint values were selected such that NMDA activation only occurred when neural activity in the compartment increased above baseline levels. To compute the overall proportion of activated NMDA receptors, we multiplied the proportion of new activation by the proportion of receptors that were both ligand-bound and available to be activated. We then added this value to the current decayed proportion of receptor activation 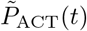 as follows (the same equation applies to both compartments):

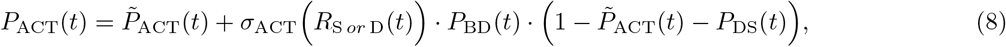

#### NMDA receptor-mediated current

Given the updated proportion of activated NMDA receptors *P*_ACT_(*t*), we then computed the NMDA receptor-mediated current to be added to each compartment (*I*_S-NMDA_(*t*) and *I*_D-NMDA_(*t*)) as a proportion of a maximum NMDA receptor-mediated current (the same equation applies to both compartments):

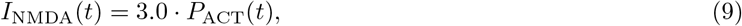

We set the maximum NMDA receptor-mediated current to 3.0, as this allowed the simulated NMDA receptor-mediated current to shape neural activity, without overtaking the influence of external inputs to the pyramidal neuron compartments.

### 4.3 BTSP weight updates

BTSP-like learning was enabled in our network at the place cell weights onto each pyramidal neuron. BTSP has been observed *in vivo* to be triggered by plateau potentials in pyramidal neurons. The weight update kernel estimated experimentally in CA1 has a notably asymmetrical shape, with a longer pre than post-plateau potential component. In addition, both components have a strong positive (potentiating) and a weaker negative (depotentiating) component [Milstein et al., 2021]. To capture these characteristics, we constructed our BTSP-like kernel (see Fig. 2D) as the product of two peak-normalized difference-of-exponential components, one rising and the other decaying:

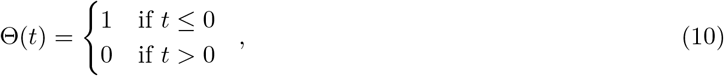

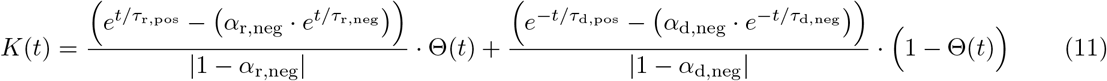

For the rising difference-of-exponentials, *τ*_r,pos_ and *τ*_r,neg_ were the time constants of the positive and negative components, respectively. For the decaying difference-of-exponentials, *τ*_d,pos_ and *τ*_d,neg_ were the time constants of the positive and negative components, respectively. Lastly, *α*_d,neg_ and *α*_r,neg_ were the weights of the negative components for the rising and decaying difference-of-exponentials, respectively.

#### BTSP kernel

To choose the parameter values, we conducted a hyperparameter search using the following targets for the BTSP kernel derived from the data and analyses reported by Milstein et al. [2021]: (1) a positive component going from -2.5 to 1.5 sec, (2) negative components returning to near-zero values at around -6 and 6 sec, and (3) a positive component peak at least 5x stronger than the negative component trough. The parameters yielded by the search are reported in Table 3.

#### BTSP weight updates

Updates to the place cell weights 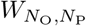 using the BTSP kernel were triggered only when a strong enough plateau potential occurred in a pyramidal neuron. We defined BTSP-triggering plateau potentials as events where a pyramidal neuron’s proximal compartment maintained a neural activity level of at least 8 (80% of the maximum, set to 10) for at least 120 ms [Epsztein et al., 2011; Bittner et al., 2015; Vaasjo et al., 2025]. Once a BTSP-triggering plateau potential occurred, this was recorded as a BTSP event and the pyramidal neuron was marked for BTSP. Given the BTSP kernel’s broad timecourse, and consistent with experimental observations [Jain et al., 2024], we simulated an additional 20 seconds (around 667 steps) after each BTSP-triggering plateau potential before applying weight updates, at which point the BTSP kernel amplitude had returned to 0. Over this time, BTSP updates for each place cell were incrementally computed online in a manner equivalent to multiplying the neural activity *R*_P_ of the input place cells (truncated to 30 sec before up to 20 sec after the plateau potential time step *t*_plateau_) with the BTSP kernel *K* (truncated to 30 sec before up to 20 sec after the kernel peak *t*_peak_). The weight updates were applied with a learning rate *λ* of 0.2 in the linear track environment and 0.15 in the open field environment (except in Fig. S1 where we assessed the effect of changing certain parameter values on place field metrics).

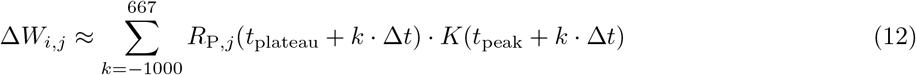

#### Divisive normalisation

To prevent runaway weight potentiation, we used divisive normalisation to penalise very high weight values. Specifically, we normalised the place cell weights of an individual pyramidal neuron *i* divisively by *β*_i_ if *β*_i_ *>* 1.0. *β*_i_ was computed as the sum of place cell weights for pyramidal neuron *i* raised to the fourth power and multiplied by a regularisation factor *α*_reg_ (2.5 for the linear track environment and 4.5 for the open field environment):

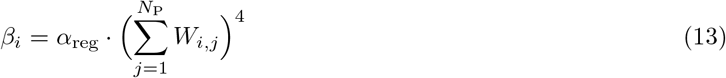

In practice, in our simulations, divisive normalisation was only triggered, i.e., the normalisation value *β*_*i*_ only surpassed 1.0, when a pyramidal neuron underwent more than one BTSP event in nearby locations. This occurred most commonly in the 2D environment simulations (see Fig. 6, S5C, S6A and S7C), but also in a few of the linear track running speed simulations (see Fig. 3B). Finally, it should be noted that we used a lower initial value for the place cell weights 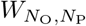 (see Table 1), as well as a weaker learning rate and a stronger regularisation factor (see Table 2) for the open field simulations compared to the linear track ones in order to compensate for the much higher number of place cells in this environment (400, compared to 60).

**Table 1.** Key circuit parameters for the linear and open field environments. The two parameters marked with an asterisk were tuned, along with the parameter marked in Table 2, using a joint hyperparameter search (see Fig. S1)

| Neuron(s) | Parameter | Linear track | Open field (if different) |
| --- | --- | --- | --- |
| All | maximum firing rate | 10 |  |
| | update time step ($\Delta t$) | 0.03 s | |
| Place (CA3) | number ($N_P$) | 60 | 400 |
|  | Gaussian field st. dev. | 0.1 m |  |
| | output weights ($W_{N_O, N_P}$) (initial) | 0.1 | 0.04 |
| Object (EC3) | Gaussian field st. dev. | 0.05 m |  |
|  | output weight(s) (fixed) | 0.4 |  |
| Inhibitory (CA1) | input filter time constant* | 0.3 s |  |
|  | input weight(s) (fixed)* | 1.0 |  |
| | sigmoid ($\sigma_S$) width | 8.0 | |
| | sigmoid ($\sigma_S$) midpoint | 6.0 | |
|  | output weight(s) (fixed) | 1.0 |  |

**Table 2.** Key pyramidal neuron (CA1) parameters for the linear and open field environments. The parameter marked with an asterisk was tuned, along with the parameters marked in Table 1, using a joint hyperparameter search (see Fig. S1)

| Category | Parameter | Compartment | Linear track | Open field (if different) |
| --- | --- | --- | --- | --- |
| general | sigmoid width | distal ($\sigma_D$) | 6.0 | |
| | | proximal ($\sigma_S$) | 8.0 | |
| | sigmoid midpoint | distal ($\sigma_D$) | 5.0 | |
| | | proximal ($\sigma_S$) | 6.0 | |
|  | maximum firing rate | both | 10 |  |
| | weight(s) (fixed) | distal → proximal | 1.0 | |
| | | proximal → distal | 0.2 | |
| NMDA | activation decay time constant | both | 0.1 sec |  |
|  | desensitisation decay time constant | both | 0.1 sec |  |
| | bound sigmoid ($\sigma_{BD}$) width | both | 20 | |
| | bound sigmoid ($\sigma_{BD}$) midpoint | both | 13.3 | |
| | activation sigmoid ($\sigma_{ACT}$) width | both | 0.5 | |
| | activation sigmoid ($\sigma_{ACT}$) midpoint | distal | 0.8 | |
|  |  | proximal | 2.0 |  |
|  | maximum current | both | 3.0 |  |
| BTSP | induction threshold | proximal | 8 |  |
|  | duration threshold | proximal | 120 ms |  |
|  | learning rate* | proximal | 0.2 | 0.15 |
| | input weight ($W_{N_P, N_O}$) regularization factor | proximal | 2.5 | 4.5 |

**Table 3.** BTSP kernel parameters.

| Parameter | Side | Component | Value |
| --- | --- | --- | --- |
| Time constant | pre (rising) | positive ($\tau_{r,pos}$) | 2.00 sec |
| | | negative ($\tau_{r,neg}$) | 2.01 sec |
| | post (decaying) | positive ($\tau_{d,pos}$) | 1.40 sec |
| | | negative ($\tau_{d,neg}$) | 1.41 sec |
| Negative weight | pre (rising) ($\alpha_{r,pos}$) | | 1.0010 |
| | post (decaying) ($\alpha_{r,neg}$) | | 1.0006 |

### 4.4 Linear track simulations

We designed the linear track as a one-dimensional, 6 meter `RatInABox` environment, with a landmark object positioned at the three-fifth mark (3.6 m from the left end of the track). On each trajectory, the agent started at the start position (0.03 m from the left end) and ran in a single direction toward to reset position (0.03 m from the right end). Once the agent was within tolerance of the reset position, the agent was returned to the start position and a new trajectory began. The tolerance for considering that the agent had reached a specific position was calculated relative to the distance the agent could cross in the next time step (0.03 s) at its current speed:

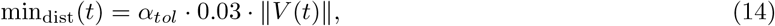

where *α*_*tol*_ is the tolerance factor and ∥*V* (*t*)∥ is the norm of the agent’s current speed vector.

#### Object visits

In the linear track simulations, a small tolerance factor *α*_*tol*_ of 0.55 was used, as the agent was only moving in one dimension, and was thus very unlikely to miss the landmark object. This meant that, at the mean speed of 0.25 m/s used in most linear track simulations, the agent had to come within 0.004 m of the landmark object for a visit to be recorded. To avoid recording a single visit as multiple different ones, a minimum of 30 steps (0.9 sec) was required between visits.

#### Agent velocity

The target mean and standard deviation of the agent’s running velocity were set using the `speed mean` and `speed std` parameters in `RatInABox`. Unless otherwise indicated, for all simulations, the target speed mean was 0.25 m/s and the target speed standard deviation was 0.05 m/s. Since the agent’s speed at each step was sampled randomly, however, the actual mean and standard deviation recorded varied a bit from these values.

#### Boundaries

To avoid boundary effects in the linear track simulations, the environment was initialised in `RatInABox` with “periodic”`boundary conditions` [George et al., 2024]. Thus, the Gaussian fields with centers near the start of the track showed some activity near the end, and vice versa. In addition, for visualisation clarity, in the simulations shown in Fig. 2 and S2, and Supp. Video 1, the agent was held for 20 sec at the start position for every trajectory except the first one. This is because, as explained above, BTSP weight updates were applied around 20 sec after a BTSP event was recorded. This holding period thus ensured that any BTSP weight updates triggered on one trajectory were applied before the agent began advancing on the trajectory. For all other simulations, the agent was not held at the start position.

#### Place field width analyses

For the place field width analyses (see Fig. 3 and S3), we assessed place field formation at 29 different running velocities, uniformly sampled between 0.05 and 0.40 m/s, inclusively. To do this, we first simulated four trajectory runs. Simulations at each speed produced exactly one BTSP event. We then simulated 15 evaluation trajectories at variable speeds (0.25 ± 0.25 m/s). In 3/29 cases, an additional BTSP event or two was recorded during the evaluation phase. These occurred within the first ten trajectories, and in each case, 15 additional evaluation trajectories were simulated after the last BTSP event had occurred. Only neural activity data from trajectories following the final BTSP weight update were used to compute the place fields, as described in the analysis section below.

#### Place field change analyses

For the analysis of place field changes (see Fig. 4 and S4), we first simulated around ten trajectories with the landmark object in the original location, at the three-fifths mark to get a robust estimate of the initial place field. We then moved the landmark object to one of 61 locations which uniformly tiled the entire track and included the original landmark object location. For the landmark object shifts that led to a second BTSP event, the event was, in each case, triggered shortly after the shift, within the first two trajectories. For all 61 simulations, we ran around six additional trajectories following the final BTSP update. Only neural activity from trajectories that occurred after the final BTSP update had been made, either at the initial object location or the final object location, were used to compute the initial, and final place fields, respectively.

### 4.5 Open field simulations

We designed two open field environment, both as two-dimensional, 2 m × 2 m `RatInABox` environments. For our first open field simulation, we aimed to test the network under conditions similar to the linear track conditions. Specifically, we aimed to test the network’s performance in a scenario where the agent typically approached the landmark object from the same direction. To encourage this, we included one interior horizontal wall extending, in (`x, y`) coordinates, from `(0, 0.4`) to `(1.2, 0.4)` (origin at the bottom left). The wall created a narrow horizontal corridor around the landmark object which we positioned toward the middle of the corridor, at `(0.5, 0.2)`. This meant that the agent was most likely to approach the object from the right (see Fig. 5A). Since the agent was moving in two dimensions, we chose a much higher tolerance (*α*_*tol*_ = 5.0) for reaching or visiting a landmark object in the open field compared to the linear track environment (see Eq. 14). As a result, moving at an average speed of 0.25 m/s, the agent would have to come within around 0.04 m of an object for a visit or target reach to be recorded.

#### Teleportation ports

We introduced teleportation into the open field corridor task as follows. We placed an entrance port in the upper area of the open field (coordinates: (1.2, 1.2)). The entrance port was left-facing, such that the agent had to be heading rightward towards the port in order to trigger a teleportation event. Specifically, the agent had to be: (1) near the entrance port (*α*_*tol*_ = 7.5), (2) on the left (within a 90^*°*^ wedge), and (3) heading toward the entrance port (also within 90^*°*^). We positioned the exit port left of the landmark object (coordinates: `(0.4, 0.2)`), and made it right-facing. When teleporting, the agent’s entry vector was reused as its exit vector, such that its exit locations were variable across teleportation events, but always toward the right, in the direction of the landmark object. This teleportation design ensured that teleportation events created variable trajectories near the landmark object, but followed a spatially consistent and predictable pattern.

In the additional teleportation simulations reported in Fig. S6, the directionality of the entrance and exit ports was flipped for half of the simulations. In these cases, the exit port was positioned to the right of the landmark object, such that the agent exited teleportation events leftward, toward the landmark object. The positions of the ports were also shifted horizontally by 0.03 to 0.1 cm in some cases to confirm that the teleportation simulation results were robust to different teleportation port positions relative to the landmark object.

#### Multiple target landmarks

For the task with multiple target landmarks, we used an open field of the same dimensions. However, this time, we randomly initialised four internal walls in the environment, each with a length between 0.2 and 0.4 m. We also initialised 40 landmark objects by randomly sampling positions uniformly across the open field. Positions were resampled if they placed a landmark object within 0.15 m of a wall or other object (see Fig. 7A).

#### Agent trajectories

In all of the open field simulations, the agent’s navigation patterns were designed to balance landmark object reaches with broad exploration of the open field to ensure good coverage for place field estimation. In the initial open field corridor simulation, therefore, we had the agent complete 12 one-minute trajectories (2000 steps each) during which it was at all times either navigating noisily toward a target landmark object or on a 9 sec (300 step) random walk. At the beginning of each trajectory, we gave the agent one of two goals: reach the target landmark object (33%) or complete a random walk (67%). If the target landmark object was reached, the random walk was completed or a new trajectory began, a new navigation goal was sampled using the same probabilities. Additional random walk periods were initiated while the agent was set to navigate toward the target landmark object if (1) the target landmark object was not in sight, i.e., it was blocked by a wall, or (2) the target landmark object had been visited within the past 30 sec (1000 steps), and no new trajectory had begun. At the end of the 12 minutes, the simulation time was extended, if necessary, until at least 10 minutes had elapsed since the weight updates had been applied for the last recorded BTSP event (see Fig. S5C). This ensured that final place fields could be reliably computed across most of the open field.

In the open field task with teleportation ports, we expanded and reweighted the navigation goals as follows: reach a target landmark object (20%), reach the teleportation entrance port (60%), and complete a random walk (20%). To simplify the interpretation of the results, teleportation events were temporarily blocked between the time when a BTSP event was recorded and when the corresponding weight updates were applied. It should be noted that the teleportation entrance port did count as reached even if the criteria for activation described above were not fulfilled. As a result, not every reach resulted in a teleportation event. The simulation was initially run for 30 minutes (60,000 steps). It was then extended until at least 6 teleportation events had occurred and at least 10 minutes had elapsed since the last BTSP weight update was applied.

Lastly, in the open field simulation with multiple target landmarks, since the target landmark objects spanned the entire environment, no trajectory resets were used. Instead, at the beginning of the simulation and each time the agent reached a target landmark object or completed a random walk, a new target was selected randomly from among the 40 landmark objects in the open field (95%) or a random walk was initiated (5%). Additional random walks were also triggered, as in the open field corridor simulations, when the target landmark object was not in sight or had already been visited within the past 30 sec (1000 steps). The simulation was run for almost 125 simulated minutes (around 250,000 steps), with no BTSP events recorded in the last 15 minutes.

#### Agent speed and direction

In 2D `RatInABox` environments, the agent’s`speed mean` parameter controls the standard deviation *σ*_Rayleigh_ of the Rayleigh distribution from which the agent’s speed is sampled at each time point [George et al., 2024]. We set this value to 0.22 m/s, and recorded linear speeds similar to those used in the linear track simulations (around 0.24 ± 0.14 m/s). During random walks, the agent’s direction of travel was sampled randomly at each time step using `RatInABox`’s agent update step with no specified direction and a `drift to random strength ratio` of 0.2. When the target object was in sight and the agent’s goal was to navigate toward it, the agent’s direction of travel was controlled using the same `drift to random strength ratio`, and a `drift velocity` *V*_drift_ computed as follows:

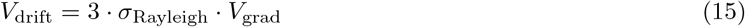

The gradient *V*_grad_ of the agent’s position with respect to its target was computed from a broad Gaussian (st. dev. of 0.6 m) centered on the target object. The same approach was used to guide the agent toward the teleportation entrance port when it was the agent’s navigation goal. As mentioned above, this did not, however, guarantee a direction of approach that would trigger a teleportation event.

### 4.6 Place field analyses

In all simulations, place fields were computed using the neural activity of the proximal compartment of each pyramidal neuron, as it represented the cell body, in addition to the radial oblique dendrites. For the linear track simulations, we computed the one-dimensional place fields of the pyramidal neurons as follows. For each period studied, e.g., before or after the landmark object was moved, only neural activity from trajectories following the last BTSP weight update was used. The neural activity recorded at each time point in the proximal compartment of the pyramidal neuron was assigned to a spatial bin, based on the agent’s position (120 spatial bins, each spanning 0.05 m). The place field was then computed as the average neural activity for each bin. For linear track simulations, place field width was calculated as the full width at half-maximum. To obtain a robust estimate, the binned neural activity was first smoothed circularly using a five-point uniform averaging kernel.

We used a similar method to compute the two-dimensional place fields of the pyramidal neurons for the open field simulations. However, in this case, we used 40×40 spatial bins, each with an area of 0.05 m × 0.05 m. Whereas in the linear track simulations, all spatial bins were typically visited by the agent during the time period used to compute each place field, this was not always the case in the open field simulations. Since no place field estimate could be obtained for spatial bins not visited by the agent, these are plotted in light grey.

### 4.7 Code

Simulations and analyses were performed in `Python 3.11` [van Rossum & Drake, 2009] and were developed using the following packages:`Matplotlib` [Hunter, 2007], `NumPy` [Harris et al., 2020], `Pandas` [McKinney, 2010], `RatInABox` [George et al., 2024], `RayTune` [Liaw et al., 2018], `SciPy` [Virtanen et al., 2020], and Seaborn [Waskom, 2021].

GitHub Copilot was used in VSCode (GitHub Copilot Chat v0.35.3) for inline suggestions to accelerate code development and facilitate consistent documentation. Suggestions were generated line-by-line, and were reviewed and edited before being incorporated. The authors take full responsibility for the content of the codebase.

## 5 Code Availability

The full codebase for running the hippocampal circuit model simulations presented in this paper and reproducing the figures is freely available on GitHub: https://github.com/colleenjg/predhpc.

## 6 Acknowledgements

The authors thank Tom George and Justin O’Hare for their valuable comments and feedback on this manuscript, and the members of the Clopath laboratory for fruitful discussions. This work was supported by Wellcome Trust 200790/Z/16/Z, Simons Foundation 564408, EPSRC EP/R035806/1, and ERC MotorAdapt 101169605.

## 7 Author Contributions

C.J.G. conceived the model. C.J.G. and C.C. designed the model. C.J.G. developed and performed the simulations. C.J.G. and C.C. wrote the manuscript.

## 10 Supplemental Videos

**Supp. Video 1: Effect of moving a target landmark object on place field shape in a linear track**. This video shows a 4 min simulation of an agent running across a linear track as in Fig. 2 at 5x speed. In this simulation, the agent completes three track traversals with the landmark object in its initial position, 3/5’s of the way down the track. The landmark object is then moved up by 0.4 m and the agent completes three more track traversals. In total, two BTSP events are triggered. The first occurs on the first track traversal, and the second occurs right after the landmark object is moved, reshaping the originally formed place field. The top panel shows in red a spatial representation of the input place cell weights to the proximal compartment of the pyramidal neuron being simulated. Following updates to the input weights, previous weights are still shown, but in a lighter shade of red. The dotted line shows the current landmark object position. Once the landmark object has been moved, a lighter dotted line shows its previous position. When applicable, the width of the input weight place field is reported in meters. Below this panel, the trajectory of the agent is represented spatially along the 6 m track. Icons show the moving agent position (black diamond), and fixed start (yellow triangle), landmark object (blue circle), and reset (red x) positions. Previous landmark object positions are shown in light blue. The bottom three panels show the neural activity of all three components of the pyramidal neuron circuit: the proximal compartment (light purple, *top*), the distal compartment (dark purple, *middle*) and the inhibitory interneuron (black, *bottom*). The x-axis shows time in simulated minutes. In the top neural activity plot, BTSP events are marked by a light purple asterisk. Landmark object visits are marked by a blue circle. In all three neural activity plots, the end of each trajectory is marked by a dashed black line.

**Supp. Video 2: Effect of teleportation on place field shape in an open field**.

This video shows the full 35 min teleportation simulation reported in Fig. 6 at 10x speed. In this simulation, the agent initially navigates either randomly or towards a landmark object. The object is located in a corridor in the open field, such that the agent will most likely approach it from the right. At each minute, the agent’s current trajectory ends and it is reset to a random location. A BTSP event is triggered the first time the agent encounters the landmark object. At 12 min, teleportation ports are added to the open field. An entrance port (“in”) is added to the upper section of the open field, and faces left. An exit port (“out”) is added left of the landmark object, and faces right, allowing the agent to approach the landmark object from the left. Once teleportation is enabled, the agent navigates either randomly, towards the landmark object or towards the teleportation entrance port. The first and second teleportation events, at around 13 and 23 min, trigger additional BTSP events, each updating the input weights to the proximal compartment of the pyramidal neuron. Four additional teleportation events occur, none of which trigger additional plasticity. The top left panel shows the open field. The landmark object is shown as a blue circle. Once teleportation is enabled, the teleportation entrance and exit ports are shown by orange triangles. Following each BTSP event, the teleportation ports are temporarily disabled, and plotted in lighter orange. The agent’s current position and head direction are shown by a circle with an arrowhead. If the agent has a current trajectory target, the target is marked by a red X. The agent is plotted in red if the target is in sight, and pink otherwise. If the agent has no current target, it is plotted in light blue. The agent’s recent positions are shown by circles that decay in size and disappear over time, and each trajectory is shown in a different colour. The top right panel shows the input place cell weights to the proximal compartment of the pyramidal neuron being simulated. The bottom three panels show the neural activity of all three components of the pyramidal neuron circuit: the proximal compartment (light purple, *top*), the distal compartment (dark purple, *middle*) and the inhibitory interneuron (black, *bottom*). The x-axis shows time in simulated minutes. In the top neural activity plot, BTSP events are marked by a light purple asterisk. Landmark object visits are marked by a blue circle and teleportation events are marked by an orange triangle and dotted line. In all three neural activity plots, the end of each trajectory is marked by a dashed black line.

**Supp. Video 3: Place field formation in a multi-target open field simulation**.

This video shows the full 124 min multi-target open field simulation reported in Fig. 7 at increasing speeds (10x, 30x, 100x, and then 300x). In this simulation, the agent navigates from landmark object to landmark object. While the agent navigates between the 40 landmark objects, the simulated pyramidal neurons undergo a total of 60 BTSP events. The top left panel shows the open field. The landmark objects are shown as blue circles. The agent’s current position and head direction are shown by a circle with an arrowhead. If the agent has a current trajectory target, the target is marked by a red X. The agent is plotted in red if the target is in sight, and pink otherwise. If the agent has no current target, it is plotted in light blue. The agent’s recent positions are shown by light blue circles that decay in size and disappear over time. The top right panel shows the maximum input place cell weights to proximal compartments across all pyramidal neurons being simulated. The bottom three panels show the neural activity of all three components of the pyramidal neuron circuit for all 40 pyramidal neurons: the proximal compartments (light purple, *top*), the distal compartments (dark purple, *middle*) and the inhibitory interneurons (black, *bottom*). The x-axis shows time in simulated minutes. In the top neural activity plot, BTSP events are marked by a light purple asterisk. Target object reaches are marked by a red X.

## 11 Supplemental Figures

**Figure S1:**
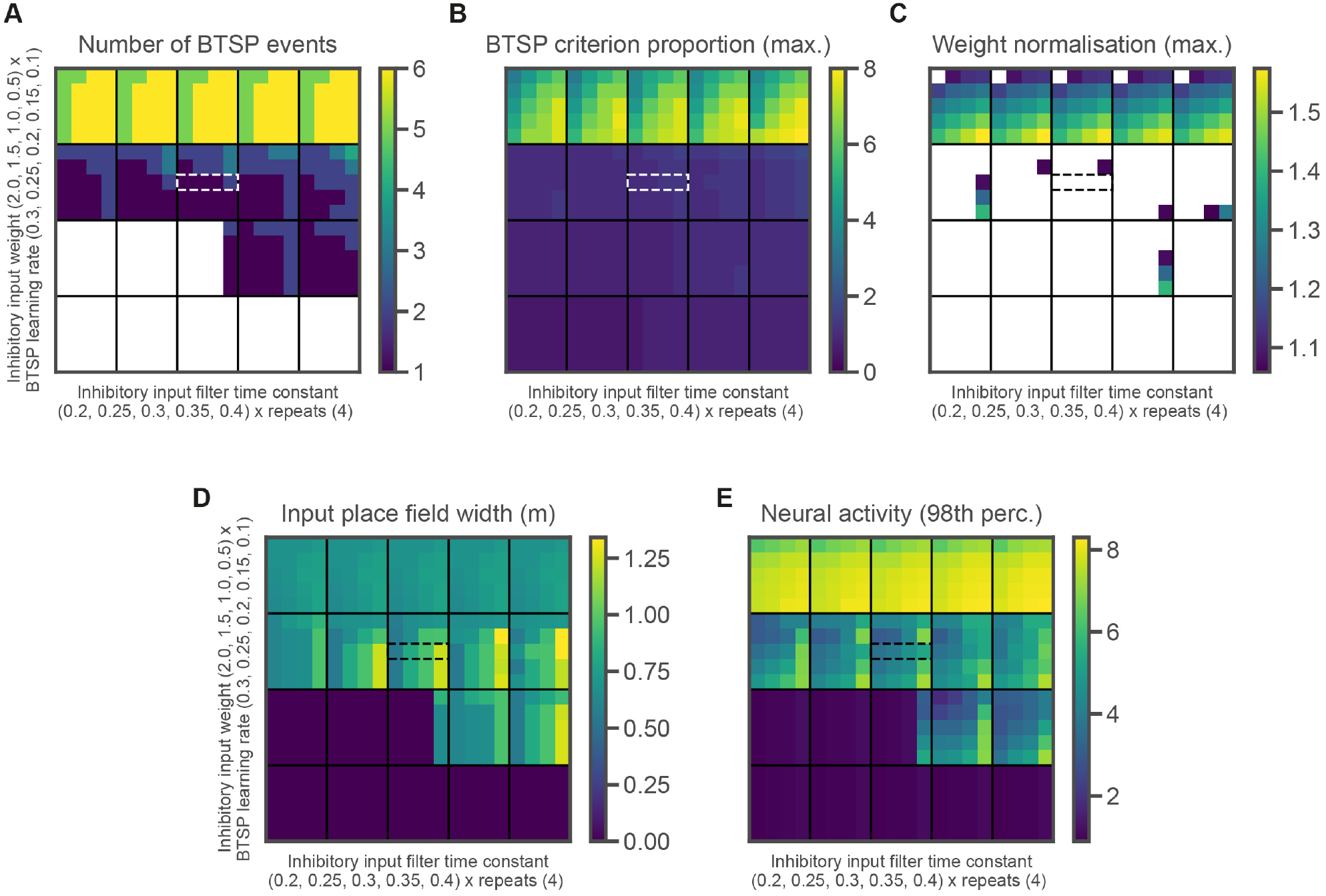
Effect of circuit parameters on linear track place field metrics. **A**. Number of BTSP events recorded during a linear track simulation (similar to Fig. 2) as a function of the values of three parameters. The simulation was run over ten linear track traversals during which the agent moved at a highly variable speed (0.25 m/s *±* 0.25 m/s). The input weight to the inhibitory interneuron is the outer variable varied along the y-axis, from top to bottom (0.5, 1.0, 1.5, 2.0). The BTSP learning rate is the inner variable varied from top to bottom within each vertical block (0.1, 0.15, 0.2, 0.25, 0.3). The time constant used to filter the input to the inhibitory interneuron is the outer variable varied along the x-axis from left to right (0.2, 0.25, 0.3, 0.35, 0.4). For each combination of parameters, the simulation was repeated four times and the results are presented within each horizontal block, sorted in ascending order from left to right. For clarity, simulations that yielded 0 BTSP events are plotted in white. The white dashed rectangle shows the final parameters combination that was selected. **B**. Similar to **A**, but showing the maximum proportion of the BTSP criterion reached across plateau potentials for each simulation. The criterion proportion increases at each consecutive time step during which the pyramidal neuron’s neural activity remains above the threshold for triggering BTSP, which is set to 8. Only criterion proportion values above or equal to 1 yield a BTSP-triggering plateau potential, reflecting at least 0.1 s spent above the neural activity threshold. Although longer plateau potentials yield higher criterion proportion values, they still only trigger one BTSP event. **C**. Similar to **A**, but showing the maximum weight normalisation applied in each simulation. The weight normalisation threshold was 1. Simulations in which no weight normalisation was applied are plotted in white. The black dashed rectangle shows the final parameter combination that was selected. **D**. Similar to **A**, but showing the final place field width (m) computed for each simulation using input place field weights. Simulations in which no place field was formed have a place field width of 0. The black dashed rectangle shows the final parameter combination that was selected. **E**. Similar to **A**, but showing the 98th percentile of the pyramidal neuron’s neural activity in each simulation. The black dashed rectangle shows the final parameter combination that was selected.

**Figure S2:**
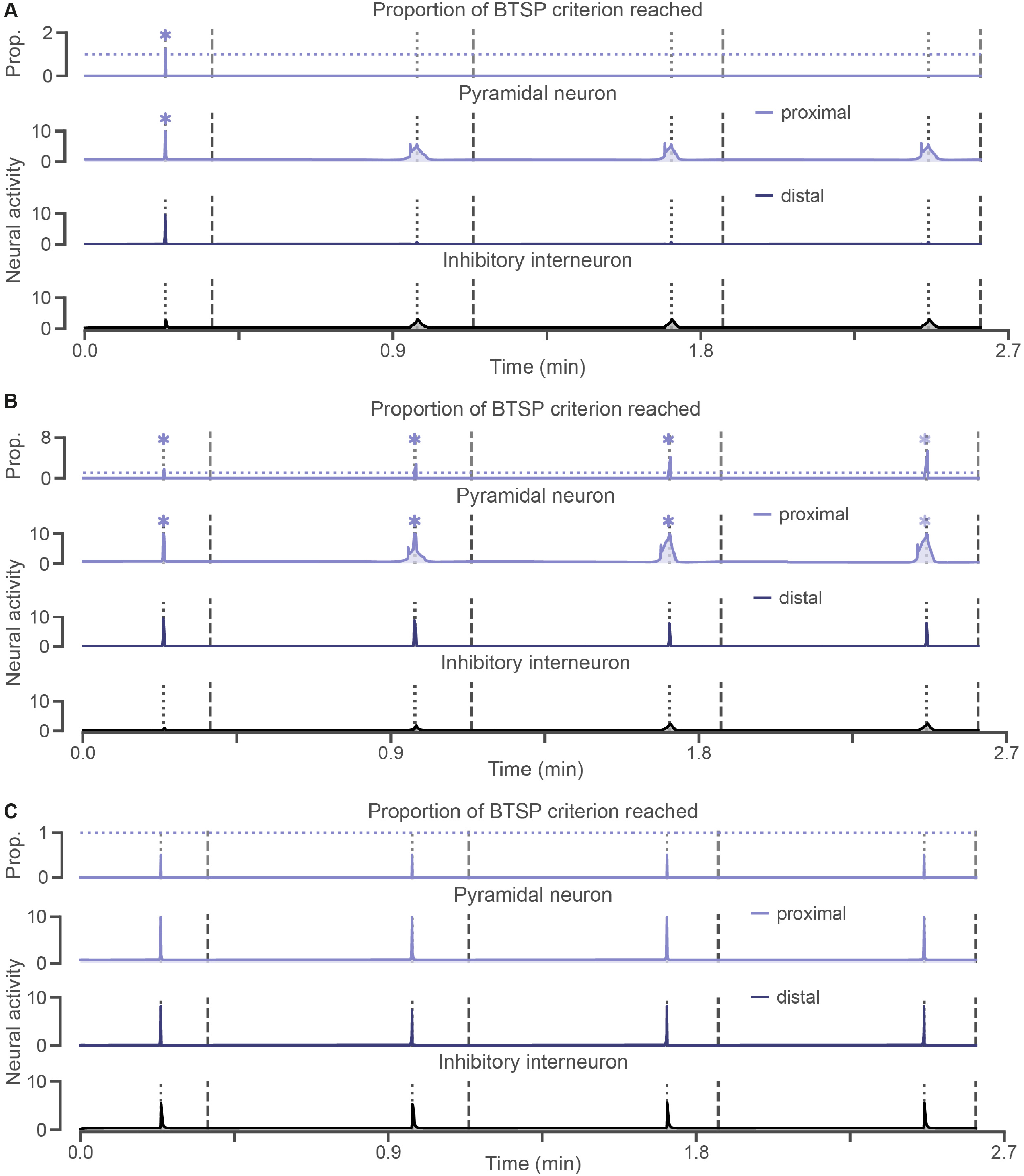
Effect of inhibitory weight on circuit self-stabilisation. **A**. Proportion of the BTSP criterion (*top*) reached at each time point (see the description in the Fig. S1B caption). Neural activity (*bottom*) across track traversals for the three components of the pyramidal neuron circuit: the proximal compartment (light purple, *top*), the distal compartment (dark purple, *middle*) and the inhibitory interneuron (black, *bottom*), as in Fig. 2A. Neural activity is expressed in arbitrary units, and ranges from 0 to 10 for each neuron or compartment. The BTSP criterion is reached and a BTSP-triggering plateau potential occurs during the first traversal only (light purple asterisk). The x-axis shows time in simulated minutes. The inhibitory weight is set to 1.0 as in Fig. 2A. (Fig. S2 caption, cont’d) **B**. Same simulation as in **A**, but with the inhibitory weight set to 0.5 instead of 1.0. The BTSP criterion is reached and a BTSP-triggering plateau potential occurs at each traversal (light purple asterisk) as distal inhibition is insufficient to prevent runaway learning. The final BTSP-triggering plateau potential is marked by an even lighter purple asterisk as the simulation ends before the weight updates it triggers can be applied. **C**. Same simulation as in **A**, but with the inhibitory weight set to 2.0 instead of 1.0. The BTSP criterion is never reached and no BTSP-triggering plateau potentials occur as distal inhibition is too strong.

**Figure S3:**
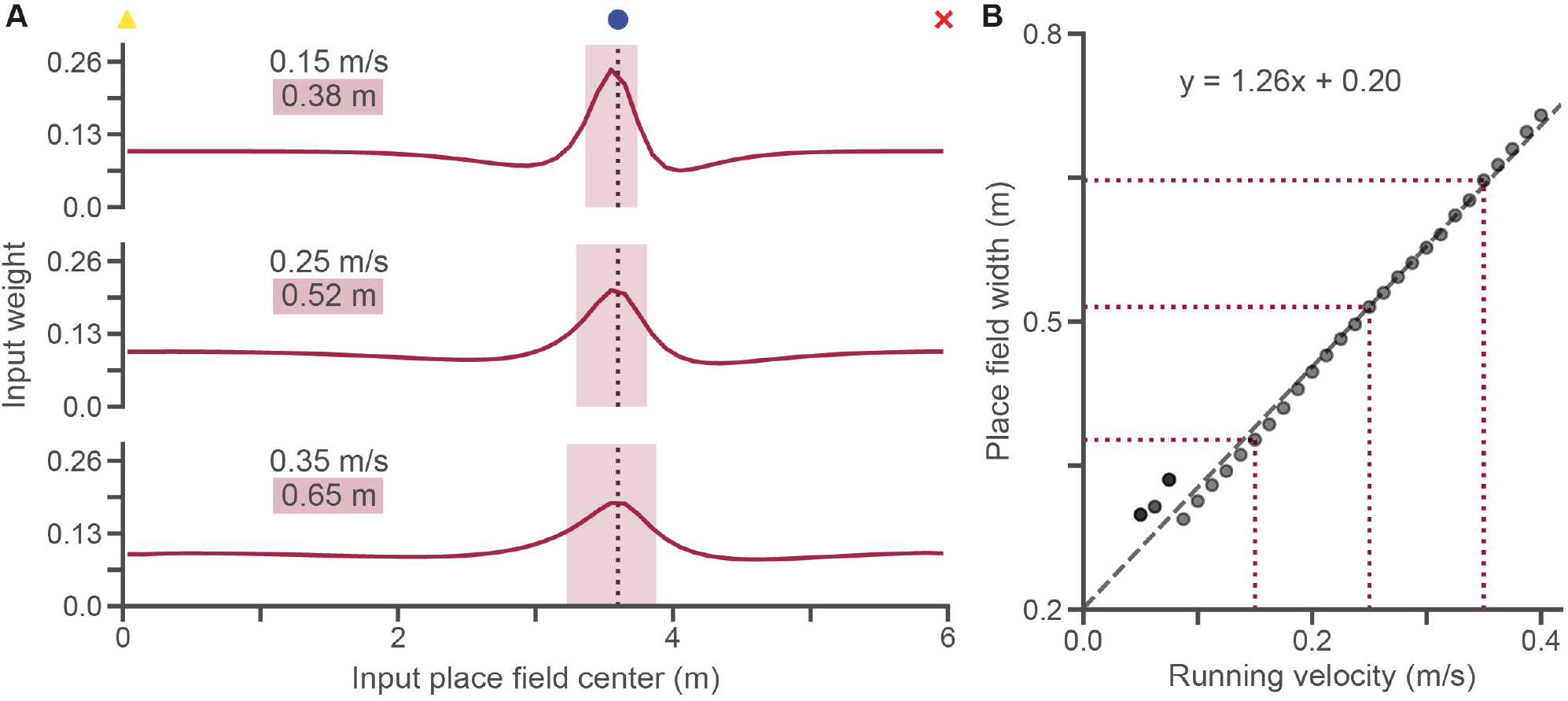
Width of input place cell weights is linearly correlated to the agent’s running velocity when BTSP was triggered. **A**. Examples of input place cell weights for different running velocities: 0.15 m/s (*top*), 0.25 m/s (*middle*), and 0.35 m/s (*bottom*), with widths: 0.38 m (*top*), 0.52 m (*middle*), and 0.65 m (*bottom*). Icons at the top show the start (yellow triangle), landmark object (blue circle), and reset (red x) positions. The vertical dashed line also shows the landmark object location. **B**. Relationship between running velocity (m/s) and place cell weight width (m). Place cell weight width increases linearly with the agent’s running velocity during the BTSP event. Data points for which a second (1/29) or third (2/29) BTSP event occurred are marked by increasingly darker dots. The dashed line shows a linear regression (*y* = 1.26*x* + 0.20). Examples from **A** are marked with red dotted lines. See Fig. 3 for the same results expressed in terms of place fields computed from the neural activity of the pyramidal neuron’s proximal compartment.

**Figure S4:**
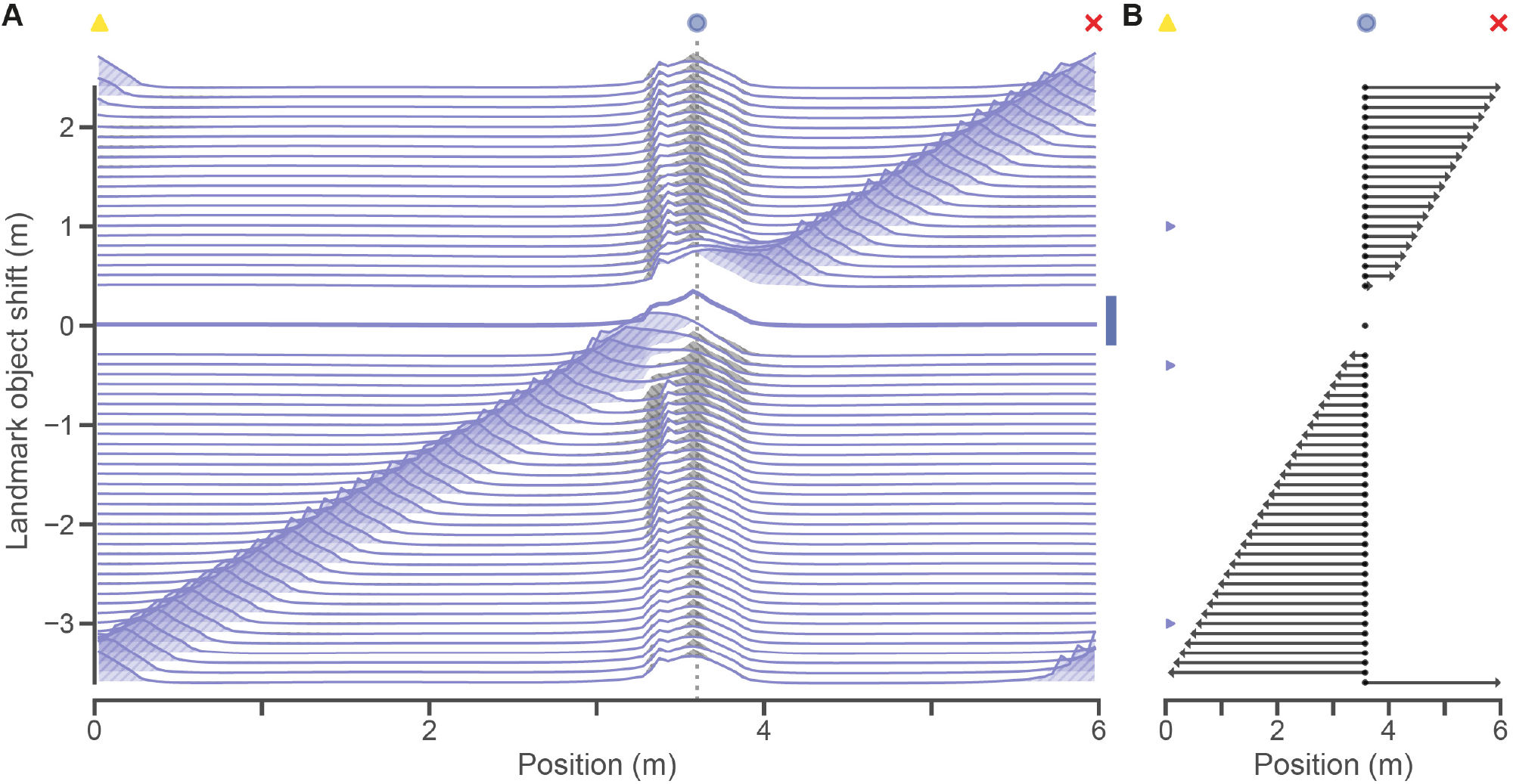
Moving the landmark object consistently leads to additional BTSP events, changing place field shapes. **A**. Place fields changed shape after the landmark object was moved. The original place field is shown at a shift of 0 (thick line) and each new place field (thinner line) is shown at its shift position along the y-axis. Light purple shading and upward hatching (//) show increases in place field amplitude, whereas grey shading and downward hatching (*\\*) show decreases in place field amplitude. Landmark shift values which did not produce a second BTSP event are left blank and marked with blue shading on the right (-0.2 and 0.3 m). Icons at the top of each subplot show the start (yellow triangle) and reset (red x) positions. The original landmark position is marked by a light blue circle and a light vertical dashed line. For clarity, the 61 shifted landmark object positions are not plotted along the x-axis. **B**. Changes in place field peak location. Icons at the top as in **A**. The examples shown in Fig. 4 are marked with light purple arrow heads.

**Figure S5:**
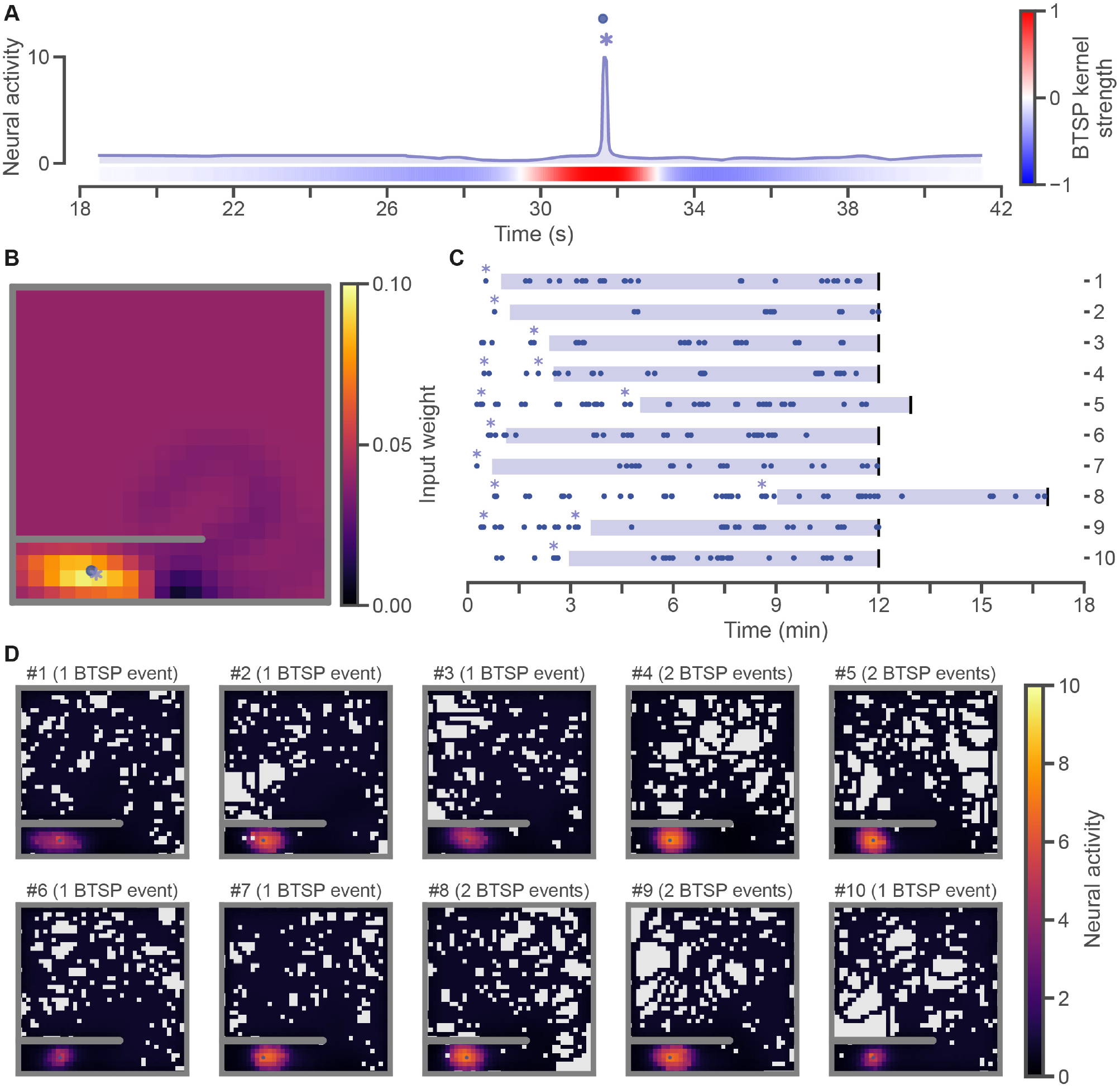
Self-stabilising place fields are consistently formed through BTSP across 2D open field simulations. **A**. Neural activity of the pyramidal neuron from Fig. 5G, zoomed in around the time of the BTSP event. Time axis is in simulated seconds. A light purple asterisk shows when the BTSP event occurred. Each visit to the landmark object is shown by a blue circle. The BTSP kernel strength is plotted as a function of time. For visual clarity, the kernel strength is plotted from -1 (most negative, blue) to 1 (most positive, red), but it should be noted that the negative and positive ranges are of different amplitudes (see Fig. 1D). **B**. Input weights from the place cells to the pyramidal neuron at the end of the open field simulation from Fig. 5. The light purple asterisk shows were the BTSP event occurred. **C**. Timelines of open field simulations run with ten different seeds. The first one is the simulation reported in Fig. 5. The time axis is in simulated minutes. Blue circles are plotted each time the agent, in each simulation, reached the landmark object. Light purple asterisks mark BTSP events. Thick vertical black lines mark the end of each simulation. Each one was run for at least 12 simulated minutes and until at least 10 minutes had passed in the simulation since the last BTSP weight updates were applied. The light purple shading shows the time span used to compute the place fields in **D**. **D**. Final pyramidal neuron place fields at the end of each simulation shown in **C**. The first place field is the same one plotted in Fig. 5E. Place fields are computed from neural activity expressed in arbitrary units ranging from 0 to 10. The number of BTSP events that occurred is reported above each simulation’s plot.

**Figure S6:**
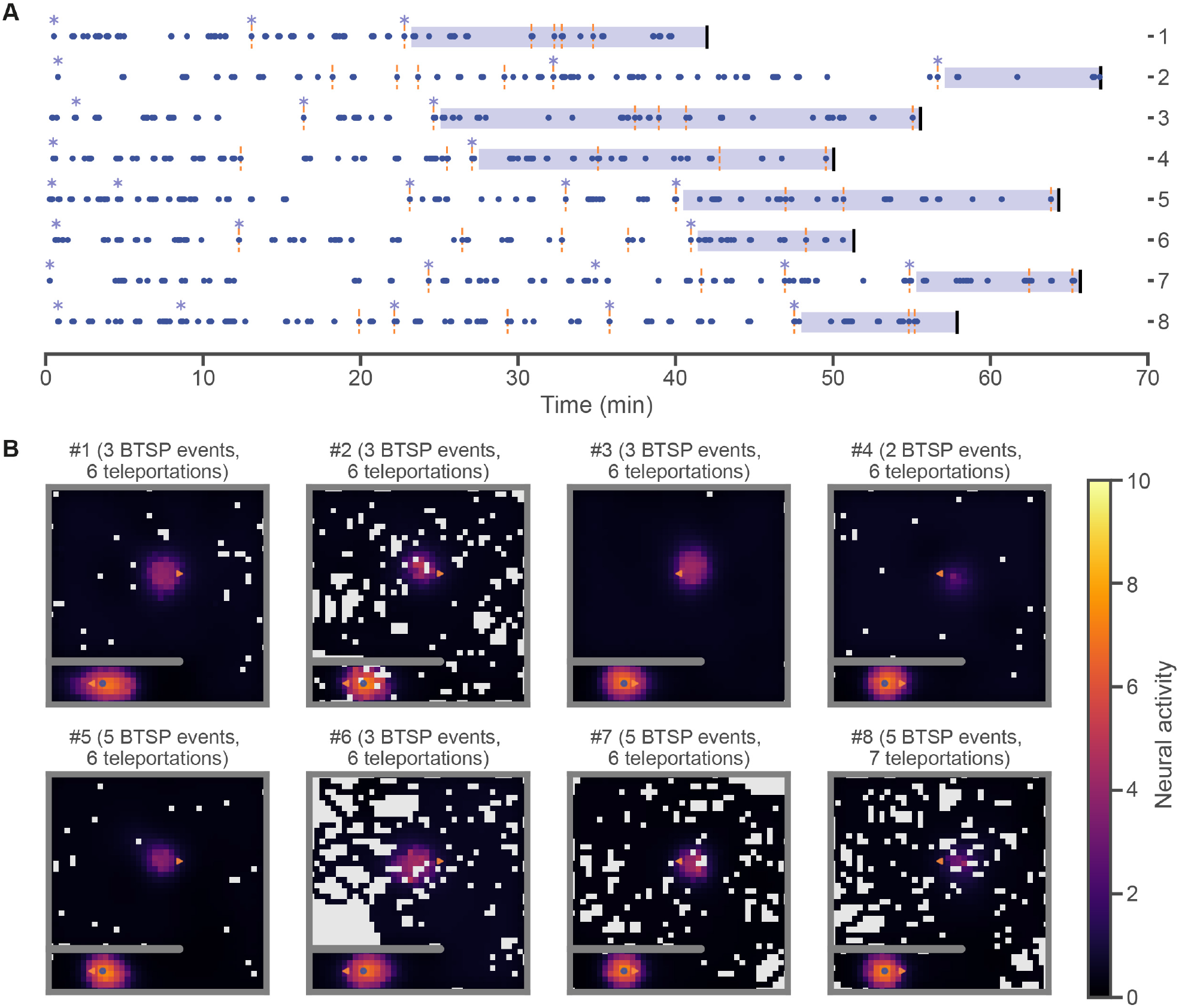
Place fields are consistently updated following teleportation events. **A**. Timelines of open field teleportation simulations run with eight different seeds. The first one is the full version of the simulation reported in Fig. 6. The time axis is in simulated minutes. Blue circles are plotted each time the agent, in each simulation, reached the landmark object. Light purple asterisks mark BTSP events. Vertical orange dashed lines mark each teleportation event. Thick vertical black lines mark the end of each simulation. Each simulation was run for at least 30 minutes and until at least 6 teleportation events had occurred and 10 minutes had passed in the simulation since the last BTSP weight updates were applied. The light purple shading shows the time span used to compute the place fields in **B**. **B**. Final pyramidal neuron place fields at the end of each simulation from **A**. The first place field is the same one as the rightmost place field plotted in Fig. 6A. Place fields are computed from neural activity expressed in arbitrary units ranging from 0 to 10. The number of BTSP events that occurred is reported above each simulation’s plot. The blue circle shows the landmark object, and the orange triangles show the teleportation ports. Teleportation ports were positioned differently across simulations. In simulations #1, #2, #5 and #6, the entrance port in the open area was left-facing, and the exit port next to the landmark object was right-facing. The opposite was the case for simulations #3, #4, #7 and #8. Furthermore, in simulations #2, #4, #6 and #8, the exit port was further away from the landmark object than in the other simulations. In most simulations, the pyramidal neuron’s place field required between one and three BTSP events following the first teleportation event to become stable. In simulations #5 and #8, however, two BTSP events occurred before the first teleportation event, and three occurred afterward before the place field became stable. In all cases, the final place field reflected the structure of the teleportation ports. Notably, the place fields formed in simulations #4 and #8 showed the weakest incorporation of the area near the teleportation entrance port. This is likely because the teleportation exit port was positioned to the right of the landmark object, and further away than in simulations #3 and #7. As a result, when the agent reached the landmark object via teleportation, its approach was much more similar to non-teleportation approaches than in other simulations.

**Figure S7:**
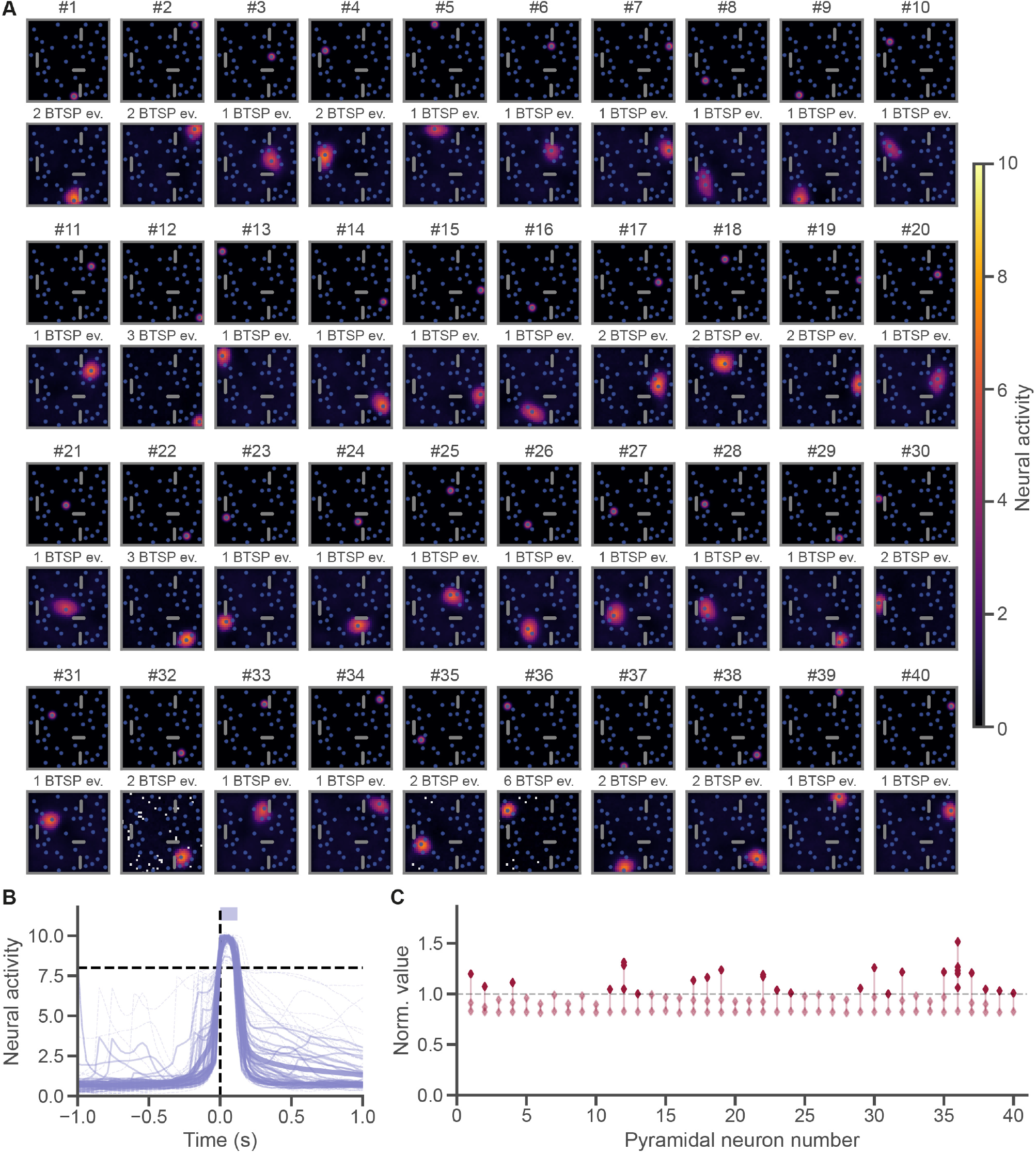
Place field formation properties across 40 pyramidal neurons with different target landmark objects. **A**. Target object cell fields (*top*) and place fields at the end of the simulation (*bottom*) for all pyramidal neurons in the multi-target open field simulation reported in Fig. 7. The number of BTSP events recorded for each of these pyramidal neurons is noted above the place field plot. Object and place fields are computed from neural activity expressed in arbitrary units ranging from 0 to 10. Place field shapes reflect the agent’s trajectory around the time of each pyramidal neuron’s BTSP events. **B**. Proximal compartment neural activity aligned to each BTSP-triggering plateau potential, overlayed for all pyramidal neurons and BTSP events. The time axis is in simulated seconds. The vertical dashed line at time 0 marks the onset of each BTSP-triggering plateau potential. The horizontal dashed line shows the minimum neural activity level (8) required to trigger a BTSP event. The light purple shading shows the minimum duration at or above the neural activity threshold required to trigger a BTSP event. **C**. Weight normalization values computed for each pyramidal neuron upon initialization and following each BTSP weight update. Vertical lines link normalization values for each neuron, chronologically. Neuron order is the same as in **A**. Only normalization values above 1.0 (dark red) are applied. Values at or below 1.0 (light red) are ignored. In most neurons, normalization was only triggered as of the second BTSP event.

